# Natural stimuli drive concerted nonlinear responses in populations of retinal ganglion cells

**DOI:** 10.1101/2023.01.10.523412

**Authors:** Dimokratis Karamanlis, Mohammad H. Khani, Helene M. Schreyer, Sören J. Zapp, Matthias Mietsch, Tim Gollisch

## Abstract

The role of the vertebrate retina in early vision is generally described by the efficient coding theory, which predicts that the retina discards spatiotemporal correlations in natural scenes. It is unclear, however, whether the predicted decorrelation in the activity of ganglion cells, the retina’s output neurons, holds under gaze shifts, which dominate the natural visual input. We here show that species-specific gaze patterns in natural stimuli can drive strong and correlated spiking responses both within and across distinct types of ganglion cells in marmoset as well as mouse retina. These concerted responses violate efficient coding and signal fixation periods with locally high spatial contrast. Finally, novel model-based analyses of ganglion cell responses to natural stimuli reveal that the observed response correlations follow from nonlinear pooling of ganglion cell inputs. Our results reveal how concerted population activity can surpass efficient coding to detect gaze-related stimulus features.

---

A prominent theory of retinal function comes from the idea of efficient coding (Barlow, 1961), which sees the retina as providing an efficient representation of incoming natural stimuli, thereby reducing the inherent spatial and temporal redundancies. This has been used to explain retinal structure and response characteristics, including center-surround receptive fields (Atick and Redlich, 1992) and the emergence of cell types and their relative spatial alignment (Karklin and Simoncelli, 2011; Ocko et al., 2018; Roy et al., 2021).

However, the decorrelation prediction of efficient coding has so far only been tested with artificial stimuli that share some statistical similarities with natural scenes (Maheswaranathan et al., 2018; Pitkow and Meister, 2012; Puchalla et al., 2005; Simmons et al., 2013), but mostly consist of static images with occasional object movement. Instead, the natural retinal input is dynamically structured by eye and head movements that rapidly shift the retinal image (Land, 2015). Large gaze shifts can induce response transients at fixation onset in neurons at the early stages of the visual system (Miura and Scanziani, 2022; Noda and Adey, 1974), and natural inputs drive nonlinear retinal processing (Heitman et al., 2016; Karamanlis et al., 2022; Shah et al., 2020; Turner and Rieke, 2016; Turner et al., 2018; Yu et al., 2022), which is currently missing from models of efficient coding. Here, we sought to study whether stimulus correlations in gaze-based natural movies are efficiently discarded by the retina.

## Natural movies can drive correlated ganglion cell responses

To test the decorrelation prediction of efficient coding, we generated natural movies for the retina (**Fig. 1A-B**) by using gaze traces from head-fixed marmosets that were viewing natural images (Yates et al., 2021). We then presented these movies to the isolated marmoset retina while recording with multielectrode arrays from retinal ganglion cells (**Fig. 1C**), which we functionally separated into the four numerically dominant types of the primate retina (**Fig. S1**): ON-, OFF-parasol and ON-, OFF-midget cells. Natural movies generated strong responses (**Fig. 1D**), which often displayed considerable correlations for pairs of neighboring cells of the same type (**Fig. 1E**). Especially ON parasol cells were strongly correlated with each other and revealed almost no decorrelation when compared to the light-intensity correlation inherent in the stimulus (**Fig. 1F**), whereas OFF midget cells, for example, displayed a high degree of decorrelation. When considering pairs of ON and OFF cells, one might expect negative response correlations, owing to their opposite preference for increases or decreases in light intensity. This was the case for midget cells, but ON and OFF parasol cells were positively correlated. These results suggest that, depending on the cell type, response decorrelation under natural stimulation can be minimal, and pairwise correlations within and between specific types of retinal ganglion cells may carry information beyond their receptive-field activation.

**Fig. 1.**
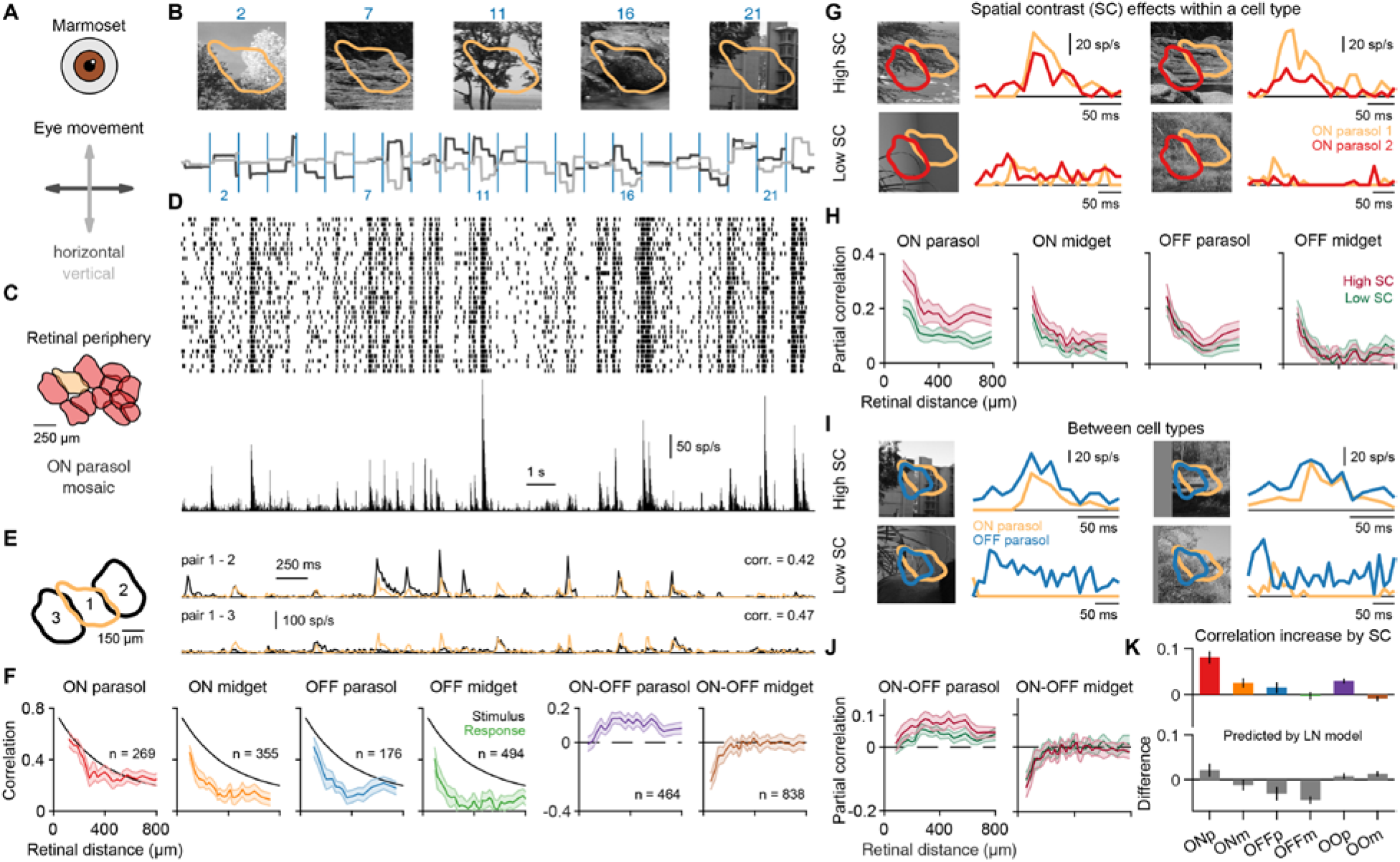
Spatial contrast in natural movies leads to concerted responses within and across ganglion cell types of primate retina. (**A**) We generated marmoset-specific movies by shifting natural images according to gaze traces recorded from head-fixed marmosets. (**B**) Each image was presented for 1 s (annotated by the blue lines) and displaced in both x- and y-directions. The receptive field of a sample ON parasol ganglion cell is overlaid. (**C**) Receptive-field mosaic of simultaneously recorded ON parasol cells. (**D**) Spike raster for 30 trials (top) and the corresponding peri-stimulus-time-histogram (bottom) for the sample ON parasol cell. (**E**) Neighboring ON parasol cells show correlated responses, quantified with Pearson’s correlation (corr.). (**F**) Correlation coefficients for ganglion cell pairs under the natural movie as a function of receptive-field distance. Colored lines represent average correlation for pairs at similar distance (with 95% confidence intervals) within the same ganglion cell type (or for pairs of ON and OFF cells) from three retinas. For reference, black lines show the correlation between stimulus pixels. (**G**) Responses of two neighboring ON parasol cells to fixations with similar light intensity, but either high (top) or low spatial contrast. (**H**) For each cell type, pairwise correlations were split into a sum of a high- and a low-spatial-contrast partial correlation. (**I**) Same as (G) for a pair of ON and OFF parasol cells. (**J**) Same as (G), but between types of different response polarity. (**K**) Median differences between high- and low-spatial-contrast partial correlations varied across types (top), and these differences diverged from differences calculated with classical linear-nonlinear (LN) models fit to cells of the same type (bottom). For ON parasol (ONp), ON midget (ONm), OFF parasol (OFFp) and ON-OFF parasol (OOp) cell pairs, the correlation increases by spatial contrast were statistically significant (p<10^-3^, Wilcoxon sign-rank test). Error bars are median ± robust confidence interval (95%).

To examine the cell-type dependence of the decorrelation, we turn to nonlinear processing within the receptive field, which can lead to cell-type-specific sensitivity to spatial contrast on spatial scales below the receptive-field size (Enroth-Cugell and Robson, 1966; Karamanlis and Gollisch, 2021; Liu et al., 2022; Turner and Rieke, 2016). Such spatial contrast, which is high when edges or textures are present in natural scenes, can particularly drive parasol cell responses in the macaque retina (Turner and Rieke, 2016; Turner et al., 2018; Yu et al., 2022). We therefore aimed at identifying whether sensitivity to spatial contrast directly influenced the pairwise response correlations. For each cell pair, we separately investigated fixation periods with high and low spatial contrast (**Fig. 1G**), while ensuring that light-intensity effects from stimulation of the linear receptive fields were balanced between the sets of high- and low-spatial-contrast fixations (see Methods). Indeed, we found that, in ON parasol cells, fixations in the high-spatial-contrast group led to stronger responses that were also more correlated than for the low-spatial-contrast fixations, in particular for shorter distances (**Fig. 1H**). These effects of high spatial contrast also existed in ON midget and OFF parasol cells, albeit to a lesser degree, but not in OFF midget cells (**Fig. 1H**). For pairs of ON and OFF parasol cells, we also observed stronger correlations for fixations in the high-spatial-contrast group relative to the low-spatial-contrast group, but not for pairs of ON and OFF midget cells (**Fig. 1I-J**). Thus, spatial stimulus structure can promote response correlations for certain cell pairs. To test whether these spatial-contrast-driven correlations arise due to standard linear processing in the retina (Pitkow and Meister, 2012), we performed the same analysis with predictions of linear-nonlinear (LN) models of ganglion cells, which capture the properties of receptive fields and spike-generation nonlinearities. The LN model failed to reproduce the observed spatial-contrast-dependent correlation differences across cell types (**Fig. 1K**), and we thus hypothesized that natural movies drive mechanisms beyond the linear receptive field that increase pairwise response correlations.

To test whether spatial-contrast-dependent correlations generalize across species, we also recorded from ganglion cells in the isolated mouse retina (**Fig. S2–4**), to which we presented natural movies generated by pairing horizontal gaze traces recorded from freely moving mice (Meyer et al., 2020) with natural images from a standard database (van Hateren and van der Schaaf, 1998). We functionally identified (**Fig. S3**) the four types of alpha ganglion cells (Baden et al., 2016; Goetz et al., 2022; Krieger et al., 2017) and, similar to the marmoset, found substantial pairwise correlations under the natural movie for certain types. We also observed spatial-contrast-dependence of pairwise correlations that was not captured by cell-type-specific LN models (**Fig. S4**). Thus, the correlation-boosting characteristics of high-spatial-contrast fixations are a general phenomenon related to nonlinear mechanisms involved in the retinal processing of natural movies.

## The nonlinear receptive field captures responses to natural images

To uncover the origin of spatial-contrast sensitivity of response correlations present in multiple ganglion cell types, we used the nonlinear subunit framework that was introduced to explain response characteristics of Y cells of the cat retina (Victor and Shapley, 1979). This framework partitions the receptive field of a ganglion cell into smaller subunits whose outputs are nonlinearly summed. These subunits are thought to correspond to bipolar cells that provide excitatory input to ganglion cells (Demb et al., 2001; Liu et al., 2017). Fitting the parameters of a nonlinear subunit model to spiking data is still an ongoing challenge (Liu et al., 2017; Maheswaranathan et al., 2018; Shah et al., 2020; Zapp et al., 2022). We therefore developed a new approach to fit subunit models to data, which we call the subunit grid model (**Fig. 2C**). Our framework is based on the simplifying assumption that individual ganglion cells receive their primary excitatory inputs from a set of identical bipolar cells that are spaced semi-regularly, thus forming a grid and performing a convolutional operation. We fit subunit grid models with a stimulus that can strongly drive cell responses, sinusoidal gratings of varying orientation and spatial frequency (**Fig. 2D**). To constrain the selection of subunits during model fitting, we introduced a density-based regularization. The resulting models succeeded in accurately capturing responses to gratings over the range of applied spatial frequencies for both marmoset and mouse (**Fig. S5–6**).

**Fig. 2.**
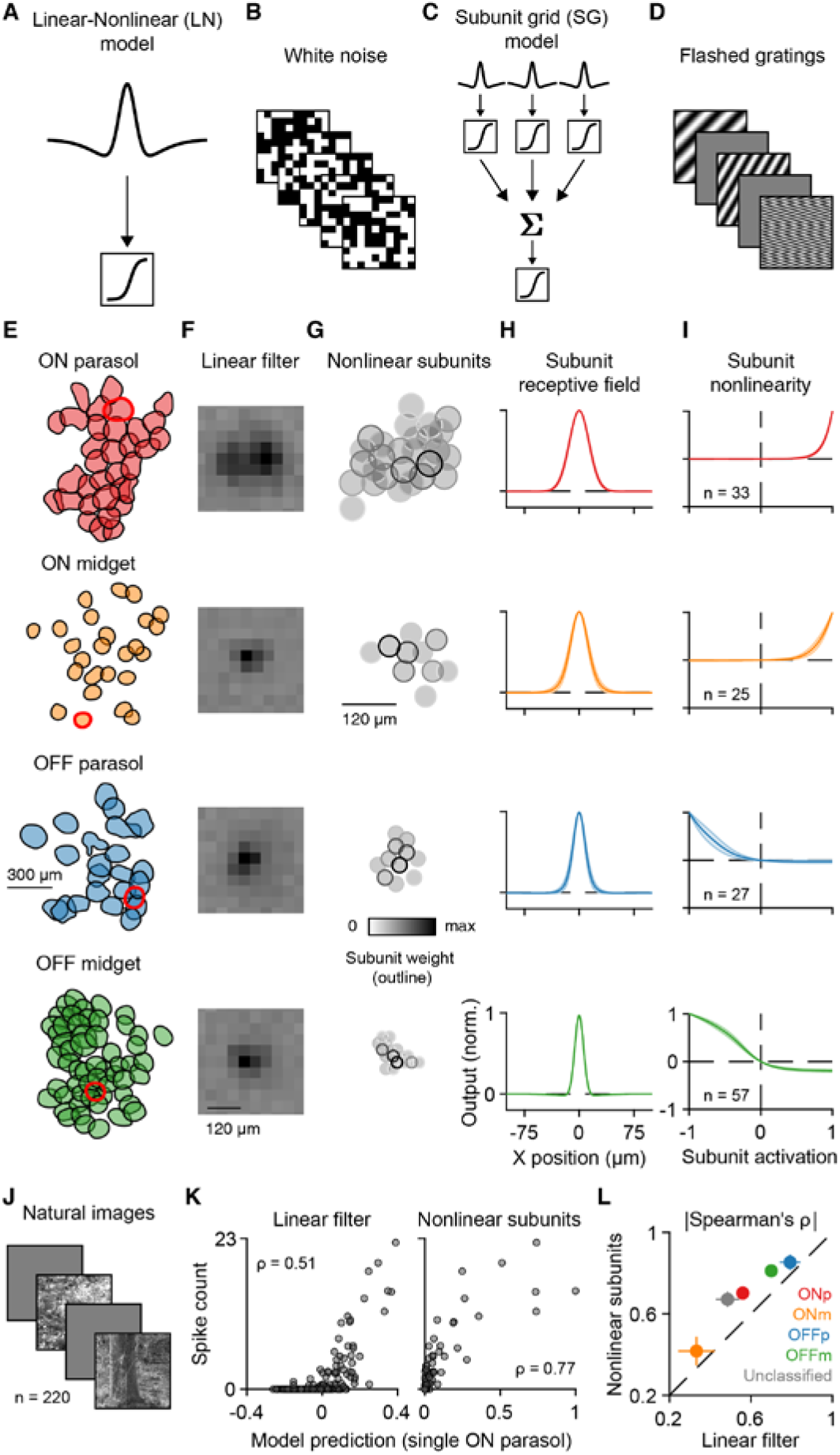
The subunit grid model captures the nonlinear receptive field and responses to natural images. (**A**) The LN model integrates light intensity over space linearly and the output is passed to a nonlinearity to generate a spike count. (**B**) We fitted LN models using responses to spatiotemporal white noise. (**C**) After light signals are processed by several identical subunits located on a hexagonal grid, the subunit grid (SG) model integrates these signals nonlinearly over the receptive field. (**D**) We fitted SG models with responses to 200-ms flashes of gratings with varying orientation, spatial frequency, and phase. (**E**) Receptive field mosaics of different ganglion cell types, with highlighted single cells (red). (**F**) The spatial filters of the highlighted cells for each type. (**G**) Obtained subunit maps for the same cells. Each circle corresponds to the 2σ Gaussian contour of the subunit center, and the shade of the circle outline corresponds to the subunit weight. (**H**) Spatial profiles of subunits, show as mean and 95% confidence interval across all cells of the same type. (**I**) Subunit nonlinearities. (**J**) Sample natural images that were flashed onto the retina for 200 ms each. (**K**) Output from the linear filter of the LN model (left) and from the summed nonlinear subunits of the SG models (right) plotted against natural image responses for a single cell. (**L**) Comparison of model performance across different types from three retinas: ON parasol (ONp, n=63), ON midget (ONm, n=31), OFF parasol (ONp, n=67), and OFF midget (OFFm, n=79) cells. Gray dot marks all cells that were unclassified (n=193), but reliable. Error bars are median ± robust confidence interval (95%).

Model fits revealed component differences between ganglion cell types (**Fig. 2E-I**). Interestingly, ON parasol cells showed more rectified nonlinearities compared to OFF, opposite of what is expected from finding in the macaque retina (Turner and Rieke, 2016). Midget cells also showed substantial rectification, consistent with findings in the peripheral macaque retina (Freedland and Rieke, 2022; Freeman et al., 2015). Cell-type-specific differences in nonlinear components were also prominent in the mouse retina (**Fig. S6**). Besides rectification in ONα types, we observed small subunits with prominent saturation in transient-OFFα cells. This nonlinearity is consistent with increased sensitivity to spatial homogeneity (Karamanlis and Gollisch, 2021) and was observed particularly for dorsal transient-OFFα cells (**Fig. S7**). Together, these results suggest that subunit grid models offer compact descriptions of nonlinear processing within the receptive fields of various ganglion cell types, including within-type differences across the retinal surface.

To connect the nonlinear receptive field to naturalistic spatial structure, we flashed natural images to the retina and recorded spike-count responses (**Fig. 2J**). We found that the derived nonlinear subunits were typically better than linear receptive fields at predicting these responses (**Fig. 2K**). These improvements in prediction performance were consistent across the numerically dominant primate cell types as well as unclassified cells (**Fig. 2L**). Improvements were also evident in the mouse retina (**Fig. S5**), with the exception of sustained-OFFα cells, which typically displayed rather linear subunit outputs, as well as transient-OFFα cells, which were well-predicted by a linear receptive field, as reported previously (Karamanlis and Gollisch, 2021). Thus, cell-type-specific models of the nonlinear receptive field are needed to capture the retinal output under naturalistic stimuli, suggesting that nonlinearities may help explain differences in spatial-contrast-driven correlations between types.

## The nonlinear receptive field predicts natural movie responses and pairwise correlations

To determine the influence of the nonlinear receptive field on pairwise correlations during natural movies, we expanded our subunit grid framework to also capture dynamic stimuli. We fitted spatiotemporal subunit grid models to ganglion cell responses under sinusoidal gratings flickering in rapid succession (**Fig. 3A**). Besides a weight map of subunits (**Fig. 3B**) and a subunit spatial profile (**Fig. 3C**), we also obtained temporal components associated with the center and the surround of all subunits (**Fig. 3D**). These components matched spatial and temporal filters derived from white-noise stimulation (**Fig. 3E**). Fitted models captured natural movie responses for different types of ganglion cells, both in the marmoset and the mouse (**Fig. 3F**). In particular, they reproduced response peaks not predicted by simple LN models. These results held for most types (**Fig. 3G**), except for mouse sustained- and transient-OFFα cells because of their nearly linear receptive fields (**Fig. S6J**). Moreover, response predictions of the subunit grid model outperformed those of alternative subunit identification schemes, such as spike-triggered non-negative matrix factorization (**Fig. S8**), originally developed for the salamander retina (Liu et al., 2017), and spike-triggered clustering (**Fig. S9**), previously applied to macaque OFF parasol cells (Shah et al., 2020).

**Fig. 3.**
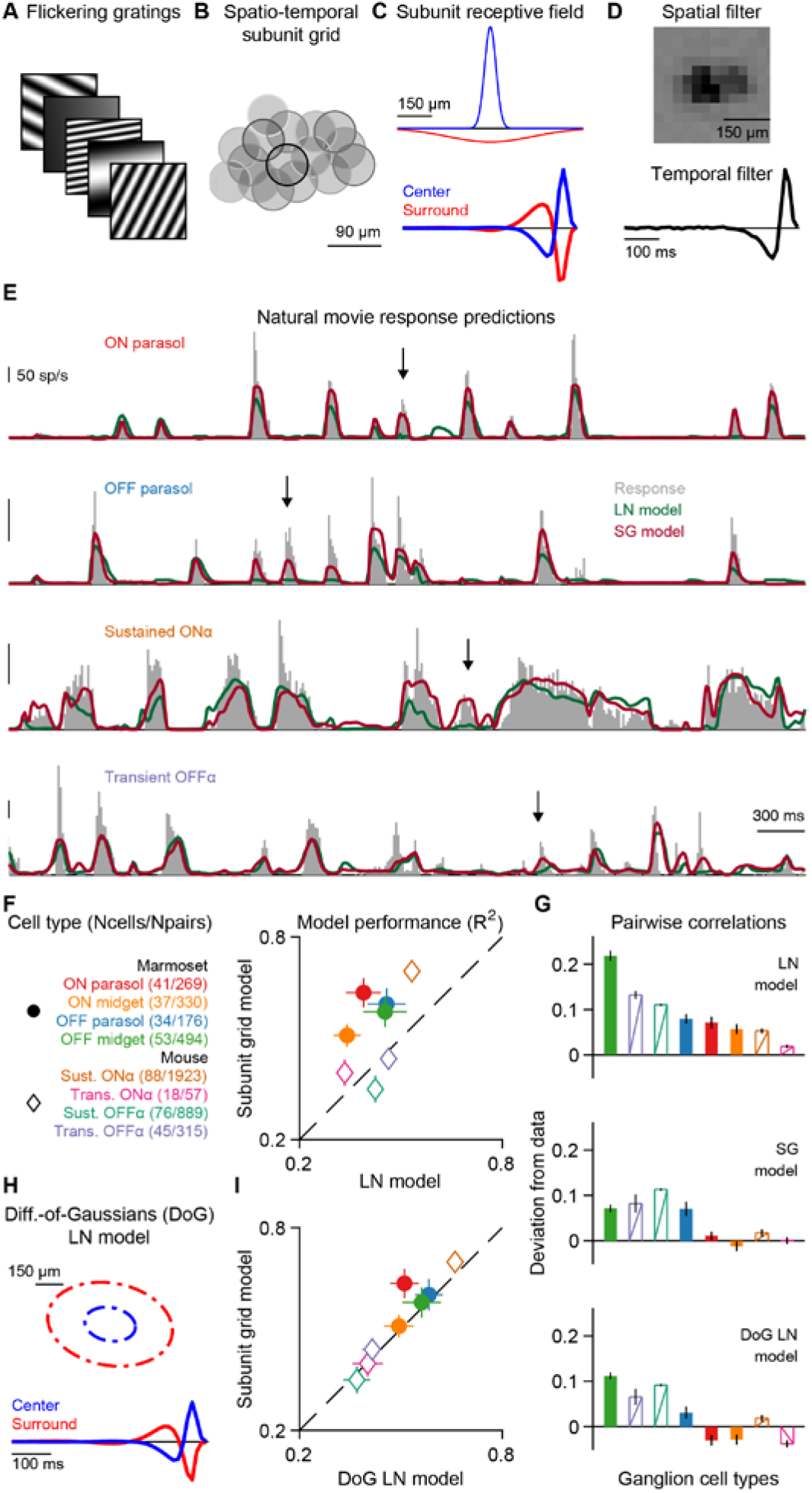
The subunit grid model captures the retinal output under natural movies. (**A**) To build spatiotemporal subunit grid models, we stimulated the retina with a rapid sequence of gratings varying in spatial frequency, phase, and orientation. (**B**) Subunit layouts for a single ON parasol cell. Each circle corresponds to the 2σ Gaussian contour of the subunit center, and the shade of the circle outline corresponds to the subunit weight. (**C**) Corresponding center and surround spatial and associated temporal components. (**D**) Linear-nonlinear (LN) model components determined with white-noise stimulation, matching the subunit layout and temporal center filter in (C). (**E**) LN and SG model predictions of ganglion cell responses to natural movies for sample cells of different types in the marmoset and the mouse retina. Arrows mark example response peaks captured by the SG but not the LN model. (**F**) We compared median model performance for all identified types in the marmoset and mouse retinas. (**G**) Deviations of pairwise correlation coefficients for model predictions from the actual correlations in the data. (**H**) Schematic depiction of DoG LN model, showing elliptical outlines of center and surround and corresponding temporal components. (**I**) Comparison of median model performance between DoG LN and SG model for different cell types. For panels (F-I), error bars are median ± 95% robust confidence interval.

The pairwise response correlations were also better captured by the subunit grid model than by the LN model (**Fig. 3H**), which tended to overestimate these correlations as previously reported (Simmons et al., 2013). That the subunit grid model predicted smaller response correlations than the LN model might seem counterintuitive, as the subunits confer sensitivity to spatial contrast, which, as we have seen, boosts correlations in the data (Fig. 1). However, the subunit grid model also captured surround suppression via the subunit surround, whereas LN models fitted to white-noise stimuli may underestimate the receptive-field surround (Wienbar and Schwartz, 2018). To distinguish the effects of captured surround suppression and spatial-contrast sensitivity conveyed by the subunits, we compared the subunit grid models to difference-of-Gaussians (DoG) LN models fitted directly to the flickering grating responses, the same stimulus as used for the subunit grid (**Fig. 3A**). The DoG LN models had surround dynamics similar to those of the subunit grid models (**Fig. 3H**), and their response predictions were better than those of white-noise-fitted LN models but still lower than the predictions of subunit grid models for certain cell types (**Fig. 3I**). Pairwise correlations estimated by DoG LN models matched those of the subunit grid models (**Fig. 3G**), confirming that surround suppression is essential for reducing the overestimation of correlations by the standard LN model and capture the correct range of response correlations. Yet, unlike the subunit grid model, the DoG LN model does not contain spatial nonlinearities that could underlie the dependence of correlations on spatial stimulus structure, as seen above (**Fig. 1**).

## Paired nonlinear responses are correlated

How does the nonlinear receptive field contribute to the response correlations observed under natural stimuli? To investigate this question, we separated the fixations for each cell pair according to how important nonlinear spatial processing was to determine the cells’ responses (**Fig. 4A**). Specifically, we tagged those fixations as nonlinear for which the predictions of the (spatially nonlinear) subunit-grid model differed most from the predictions of the (spatially linear) DoG LN model. For these “maximally-differentiating fixations” (here the 20% of fixations with the largest prediction differences), the subunit grid model displayed superior model predictions compared to the DoG LN models for nonlinear cell types, such as ON and OFF parasols in the marmoset and sustained-ONα cells in the mouse (**Fig. 4B**). Nonlinear receptive field activation was also highly relevant for pairs of parasol cells of opposite polarity, but not for midget cells or pairs of mouse ON and OFF cell types (**Fig. 4B**). Thus, the subunit grid model can indeed capture responses particularly well when nonlinear spatial processing becomes important.

**Fig. 4.**
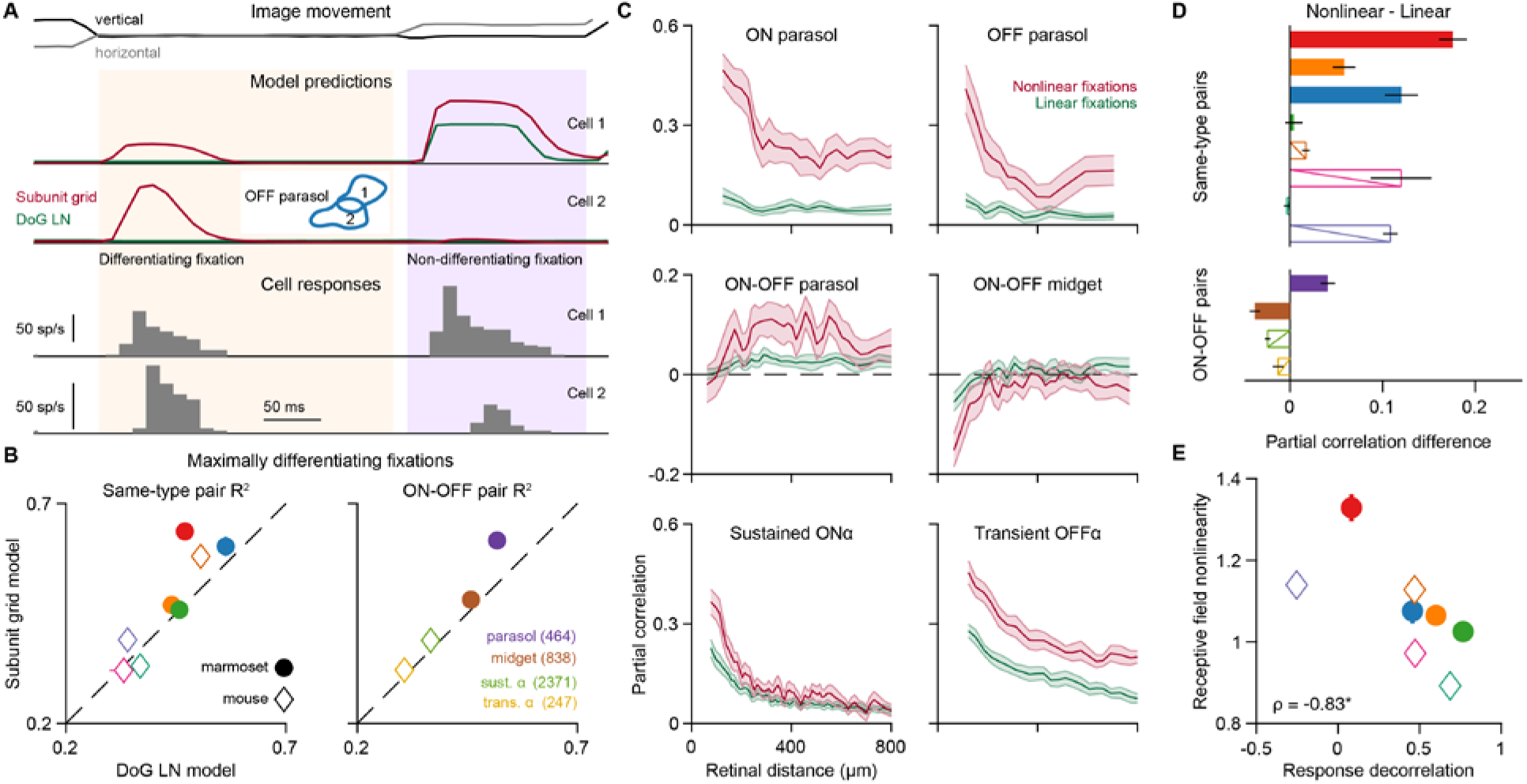
Concerted responses during fixations evoking nonlinear responses. (**A**) Image displacement traces during the presentation of a single image. Model predictions (middle) for two neighboring OFF parasol cells (receptive fields in inset) for the DoG LN and the subunit grid models. Shaded areas mark two consecutive fixations. Because model predictions strongly differ during the first fixation across both cells, the first fixation is a differentiating fixation and the second one a non-differentiating fixation. Responses (bottom) of the two cells during the same period. (**B**) Comparison of average model performances (R^2^) for the subunit grid model and the DoG LN model during the top 20% maximally differentiating fixations. Color scheme and cell pair numbers for the left plot match Fig. 3F. (**C**) Contributions of linear and nonlinear fixations to the total pairwise correlations determined for the movie (with 95% confidence intervals). (**D**) Summary of correlation differences in (E) across multiple ganglion cell types in the mouse and marmoset. (**E**) The relationship between receptive field nonlinearity, calculated as the performance improvement during maximally-differentiating fixations, and overall pairwise response decorrelation, calculated as the sign-inverted decrease of pairwise correlations relative to the stimulus correlation. Star denotes a significant Spearman’s correlation (p = 0.015). For panels (B), (D), and (E), error bars are median ± 95% robust confidence interval.

The cell types for which capturing nonlinear processing via the subunit grid model was particularly relevant during maximally-differentiating fixations were also the ones that had shown a strong dependence on spatial stimulus structure in the pairwise response correlation (**Fig. 1K**). Thus, to determine whether the nonlinear receptive field alone could lead to this difference in response correlations, we divided fixations into those where responses rather followed the linear receptive field and those where responses depended on nonlinear mechanisms (based, again, on the difference of the two model predictions). We then separated the contributions of the linear and nonlinear fixations to the overall correlations by calculating partial correlations for each set. Indeed, for parasol cells in the marmoset and nonlinear alpha cells in the mouse, response correlations during nonlinear fixations contributed the most to the overall pairwise correlations (**Fig. 4C-D**). By contrast, linear cells, such as marmoset OFF midget and mouse sustained OFFα, displayed more balanced partial correlations between linear and nonlinear fixations. Moreover, nonlinear fixations were responsible for the surprising positive correlations of ON and OFF parasol cells (**Fig. 4C-D**). We thus conclude that the nonlinear receptive field drives concerted cell responses during fixations that contain salient spatial structure.

## Discussion

We provide direct evidence that classical efficient coding in the retina is violated in a cell-type-specific manner under natural stimuli that include gaze dynamics. Although some ganglion cell types displayed substantial decorrelation in line with efficient coding and redundancy reduction, other ganglion cells revealed substantially correlated activity both within and across ganglion cell types. The correlations were particularly pronounced when the stimulus shifted to a new fixation that contained high spatial contrast. This concerted activity originated in nonlinear processing within the receptive fields of retinal ganglion cells, a mechanism that was previously associated with decorrelation in the context of artificial stimuli (Maheswaranathan et al., 2018). Furthermore, the response correlations persisted, even though our stimuli contained measured fixational eye movements, which are thought to contribute to decorrelation of retinal responses (Segal et al., 2015). Comparisons of cell types revealed a unifying relation that holds across both marmoset and mouse: ganglion cells with stronger nonlinear processing tend to perform less stimulus decorrelation (**Fig. 4E**).

Because retinal circuit nonlinearities have been associated with computations underlying visual feature detection (Gollisch and Meister, 2010; Kerschensteiner, 2022), we hypothesize that the response correlations in nonlinear cell types aid in signaling the detection of a relevant visual feature in natural scenes (Lettvin et al., 1959). For example, mammalian direction-selective retinal ganglion cells (Sabbah et al., 2017), which are a prime example of feature detectors, were found to have a strongly nonlinear receptive field (**Fig. S10A-F**) and their pronounced pairwise response correlations in our data even exceeded stimulus correlations when detecting motion in their preferred direction (**Fig. S10G-J**). It seems likely that the correlated responses enforce the message that is sent to downstream brain areas about the detected feature.

This may be particularly important for distinguishing activity evoked by the relevant feature, such as local spatial contrast or the preferred motion signal, from activity that simply results from preferred mean light intensity inside the receptive field (Kühn and Gollisch, 2019) and may thus help in disentangling an otherwise multiplexed neural code (Meister et al., 1995). Moreover, the relative activity differences within coordinated spiking events may serve as a robust code for feature detection when cells with different feature preferences are considered (Kühn and Gollisch, 2019). While the occurrence of spikes in a neuron or the absolute spike count are confounded by light intensity and contrast information, or even other stimulus dimensions to which the neuron might be sensitive, such contributions may at least partially cancel out when encoding the occurrence and strength in activity differences of pairs or groups of neurons.

Efficient coding is often considered a natural assumption for sensory systems because of the need to preserve energy associated with neuronal activity (Balasubramanian and Sterling, 2009). For feature detection, however, energy consumption might not be such a fundamental constraint if the visual feature in question is rare in natural visual inputs. Then, the retinal code may multiplex correlated nonlinear responses containing feature information with decorrelated baseline activity (Deny et al., 2017). We thus argue that the retinal output can remain efficient while being robust for feature detection to allow, for instance, the occasional quick escape from a predator (Kim et al., 2020).

## Materials and Methods

### Tissue preparation and electrophysiology

We obtained retinal tissue from three adult marmoset monkeys (*Callithrix jacchus*) of male sex, aged 12, 13, and 18 years. Retinal tissue was obtained immediately after euthanasia from animals used by other researchers, in accordance with national and institutional guidelines and as approved by the institutional animal care committee of the German Primate Center and by the responsible regional government office (Niedersächsisches Landesamt für Verbraucherschutz und Lebensmittelsicherheit, permit number 33.19-42502-04-20/3458). After enucleation, the eyes were dissected under room light, and the cornea, lens, and vitreous humor were carefully removed. The resulting eyecups were then transferred into a light-tight container containing oxygenated (95% O_2_ and 5% CO_2_) Ames’ medium (Sigma-Aldrich, Munich, Germany), supplemented with 4 mM D-glucose (Carl Roth), and buffered with 20-22 mM NaHCO3 (Merck Millipore) to maintain a pH of 7.4. The container was slowly heated to 33°C, and after at least an hour of dark adaptation, the eyecups were dissected into smaller pieces. All retina pieces used in this study came from the peripheral retina (7-10 mm distance to the fovea). The retina was separated from the pigment epithelium just before the start of each recording.

We also obtained retinal tissue from eight adult wild-type mice of female sex (C57BL/6J), mostly between 7-15 weeks old (except for one 23-week-old). All mice were housed in a 12-hour light/dark cycle. Experimental procedures were in accordance with national and institutional guidelines and approved by the institutional animal care committee of the University Medical Center Göttingen, Germany. We cut the globes along the ora serrata, removing the cornea, lens, and vitreous humor. The resulting eyecups were hemisected to allow two separate recordings. Based on anatomical landmarks, we performed the cut along the midline and marked dorsal and ventral eyecups. Before the start of each recording, we isolated retina pieces from the sclera and pigment epithelium.

We placed retinal pieces ganglion cell-side-down on planar multielectrode arrays (Multichannel Systems; 252 electrodes; 10- or 30-μm diameter, either 60- or 100-μm minimal electrode distance) with the help of a semipermeable dialysis membrane (Spectra Por), stretched across a circular plastic holder (removed before the recording). The arrays were coated with poly-D-lysine (Merck Millipore). For some marmoset recordings, we used a 60-electrode perforated MEA system (Schreyer and Gollisch, 2021). Dissection and mounting were performed under infrared light on a stereo microscope equipped with night-vision goggles. Throughout the recording, retinal pieces were continuously superfused with oxygenated Ames’ solution flowing at 8-9 ml/min for the marmoset or 5-6 ml/min for the mouse retina. The bath solution was heated to a constant temperature of 33°C-35°C via an inline heater in the perfusion line and a heating element below the array.

Extracellular voltage signals were amplified, bandpass filtered between 300 Hz and 5 kHz, and digitized at 25 kHz sampling rate. We used Kilosort (Pachitariu et al., 2016) for spike-sorting. To ease manual curation, we implemented a channel selection step from Kilosort2 in the pipeline, by discarding channels that contained very few threshold crossings and probably no spiking activity (modified version available at https://github.com/dimokaramanlis/KiloSortMEA). We curated the output of Kilosort through phy, a graphical user interface for visualization, and selected only well-separated units with clear refractory periods in the autocorrelograms. In a few cases, we had to merge units with temporally misaligned templates; we aligned the spike times by finding the optimal shift through the cross-correlation of the misaligned templates.

### Visual stimulation

Visual stimuli were generated and controlled through custom-made software, based on Visual C++ and OpenGL. Different stimuli were presented sequentially to the retina through a gamma-corrected monochromatic white OLED monitor (eMagin) with 800 x 600 square pixels and 85 (marmoset) or 75 Hz (mouse) refresh rate. The monitor image was projected through a telecentric lens (Edmund Optics) onto the photoreceptor layer of the retina, and each pixel’s side measured 7.5 μm on the retina. For some marmoset recordings, the image of the OLED screen was combined with the light path of an upright microscope through a beam splitter and focused through a custom-made optics system and the 4x objective of the microscope onto the photoreceptor layer, with each pixel’s side measuring 2.5 μm on the retina. All stimuli were presented on a background of low photopic light levels, and their mean intensity was always equal to the background. For the marmoset, the background light intensity resulted in ~3000 M*/cone/s and for the mouse in ~4000 R*/rod/s. We fine-tuned the focus of stimuli on the photoreceptor layer before the start of each experiment by visual monitoring through a light microscope and by inspection of spiking responses to contrast-reversing gratings with a bar width of 30 μm.

### Receptive field characterization

To characterize spatial and temporal response properties of the recorded ganglion cells, we used a spatiotemporal binary white-noise stimulus (100% contrast) consisting of a checkerboard layout with flickering squares, ranging from 15 to 37.5 μm on the side. The stimulus update rate ranged from 21.25 to 85 Hz. We then calculated spike-triggered averages (STAs) over a 500 ms time window, and extracted spatial and temporal filters for each cell as previously described (Rhoades et al., 2019). Briefly, the temporal filter was calculated from the average of STA elements whose absolute peak intensity exceeded 4.5 robust standard deviations of all elements. The robust standard deviation of a sample is defined as 1.4826 times the median absolute deviation of all elements, which gives the correct standard deviation for a normal distribution. The spatial filter was obtained by projecting the spatiotemporal STA on the temporal filter. We also calculated spike-train autocorrelation functions under white noise, using a discretization of 0.5 ms. For plotting and subsequent analyses, all autocorrelations were normalized to unit sum.

For each cell, a contour was used to summarize the spatial receptive field (RF). We upsampled the spatial RF to single-pixel resolution, and then blurred it with a circular Gaussian of σ = 4 pixels. We extracted RF contours using MATLAB’s “contourc” function at 25% of the maximum value in the blurred filter. In some cases, noisy STAs would cause the contour to contain points that laid further away from the actual spatial RF. Thus, we triaged the contour points, and removed points that exceeded 20 robust standard deviations of all distances between neighbors of the points that were used to define the contour. This process typically resulted in a single continuous area without holes. The center of each RF was defined as the median of all contour points, and its area as the area enclosed by the contour.

### Ganglion cell type identification

We used responses to a barcode stimulus (Drinnenberg et al., 2018) to cluster cells into functional types within each single recording. This stimulus is a one-dimensional variation of light intensity that moves across the screen. In particular, the barcode pattern had a length of 12,750 (or 12,495) mm and was generated by superimposing sinusoids of different spatial frequencies (f) with a 1/f weighting. The constituent sinusoids had spatial frequencies between 1/12750 (or 1/12495) and 1/120 μm^-1^ (separated by 1/12750 or 1/12495 μm^-1^ steps, respectively) and had pseudorandom phases. The final barcode pattern was normalized so that the brightest (and dimmest) values corresponded to 100% (and −100%) Weber contrast from the background. The pattern moved horizontally across the screen at a constant speed of 1275 (or 1125) μm/s, and the stimulus was repeated 10 to 20 times. Obtained spike trains were converted into firing rates using 20-ms time bins, and Gaussian smoothing with a σ = 20 ms. We quantified cell reliability with a symmetrized coefficient of determination (R^2^), as described previously (Karamanlis and Gollisch, 2021). We only included cells with a symmetrized R^2^ value of at least 0.1 that were not putative direction-selective cells (see below).

We used average responses to the barcode stimulus to generate a pairwise similarity matrix, as described previously (Drinnenberg et al., 2018). We defined the similarity between each pair of cells as the peak of the normalized cross-correlation function between the spike rate profiles of the two cells. To obtain a final similarity matrix, we multiplied the barcode similarity matrix with three more similarity matrices, obtained from RF response properties. The first two were generated by computing pairwise correlations between both the temporal filters and the autocorrelation functions of each cell. The third one used RF areas and was defined as the ratio of the minimum of the two areas over their maximum (Jaccard index).

We converted the combined similarity matrix to a distance matrix by subtracting each entry from unity. We then computed a hierarchical cluster tree with MATLAB’s “linkage” function, using the largest distance between cells from two clusters as a measure for cluster distance (complete linkage). The tree was used to generate 20 to 50 clusters; we chose the number depending on the number of recorded cells. This procedure yielded clusters with uniform temporal components and autocorrelations, and RF overlaps expected from tiling, but typically resulted in oversplitting functional ganglion cell types. Thus, we manually merged clusters with at least two cells, based on the similarity of properties used for clustering and on RF tiling. To incorporate cells that were left out of the clustering because of the barcode quality criterion, we expanded the clusters obtained after merging. For each unclustered cell, we calculated Mahalanobis distances to all the obtained clusters. A cell was assigned to a cluster if its Mahalanobis distance from the center of the cluster was at most 5 but at least 10 for all other clusters. Our method could consistently identify types with tiling RFs, such as parasol/midget cells in the marmoset and alpha cells in the mouse retina, which are the ones primarily analyzed in this work.

### Matching cell types to previously identified functional ganglion cell types

We validated the consistency of cell type classification by examining cell responses to a chirp stimulus (Baden et al., 2016), which was not used for cell clustering. Light-intensity values of the chirp stimulus ranged from complete darkness to the maximum brightness of our OLED screen. The stimulus was presented 10-20 times. For the mouse retina the parameters of the chirp stimulus matched the original description, which allowed us to compare cell responses to calcium traces in a database of classified retinal ganglion cells (Baden et al., 2016). To convert spikes to calcium, we convolved our spiking data with the calcium kernel reported in the original paper. We then computed correlations to the average traces of each cluster in the database.

For some mouse experiments, we used the responses under spot stimuli (Goetz et al., 2022). Briefly, we flashed one-second-long spots over the retina at different locations and with five different spot diameters (100, 240, 480, 960 and 1200 μm). Between spot presentations, illumination was set to complete darkness, and the spots had an intensity of 100-200 R*/rod/s. For each cell, we estimated a response center, by identifying which presented spot location yielded the strongest responses, combining all five spot sizes. We only used cells whose estimated response center for the spots lay no further than 75 μm from the RF center. To calculate similarities to the available database (Goetz et al., 2022), we concatenated firing rate responses to the five spot sizes and then used correlation to match our recorded ganglion cells to the database templates.

In mouse experiments, we also used saccade-like shifted gratings to detect image-recurrence-sensitive cells as previously described (Karamanlis and Gollisch, 2021; Krishnamoorthy et al., 2017). These cells correspond to the transient-OFFα cells in the mouse retina.

### Natural movies, LN model predictions, and response correlations

For the marmoset retina, we constructed natural movies based on rationale previously applied for the macaque retina (Heitman et al., 2016; Shah et al., 2020). Briefly, the movies consisted of 347 images that were shown for one second each and jittered according to measurements of eye movements obtained from awake, head-fixed marmoset monkeys (J.L. Yates, personal communication). The procedure for recording eye movements has been previously described (Yates et al., 2021). The eye movement data were collected at a scale of about 1.6 arcmin/pixel, which approximately corresponds to 2.67 μm on the marmoset retina, using a retinal magnification factor of 100 μm/deg (Troilo et al., 1993). We presented natural movies at a comparable scale on the retina (2.5 μm/pixel). We resampled the original 1000-Hz gaze traces to produce a movie with a refresh rate of 85 Hz. The presented natural movie consisted of 30-35 cycles of varying training and repeated test stimuli. Test stimuli consisted of 22 distinct natural images that matched the grayscale images viewed by the marmosets during eye movement tracing. Each test image was paired with a unique movement trajectory that also matched the marmoset eye movements. For each training stimulus cycle, we presented 40 images out of the 325 remaining images (sampled with replacement), each paired with a unique movement trajectory. These 325 images were obtained from the Van Hateren database (van Hateren and van der Schaaf, 1998) and were multiplicatively scaled to have the same mean intensity as the background. To extract firing rates for the test stimuli, we binned spike trains at a single frame resolution, and we only used cells with a symmetrized R^2^ of at least 0.2 for subsequent analyses.

For the mouse retina, we applied a similar procedure. Briefly, the movies consisted of the aforementioned 325 images from the Van Hateren database, shown for one second each and jittered according to the horizontal gaze component (Meyer et al., 2020) of freely moving mice (Arne Meyer, personal communication). We resampled the original 60-Hz gaze traces to produce a movie with a refresh rate of 75 Hz. For our recordings, the one-dimensional gaze trajectory was randomly assigned to one of four orientations (0, 45, 90 or 135 degrees) for each one-second image presentation. The amplitude of the original movement was transformed into micrometer on the retina using a retinal magnification factor of 31 μm/deg for the mouse. All images were multiplicatively scaled to have the same mean intensity as the background. Testing stimuli consisted of 25 distinct natural images, paired with unique movement trajectories. The training stimuli consisted of 35 images out of the remaining 300 (sampled with replacement), each paired with a unique movement trajectory.

All model predictions for natural movies used the test stimulus part for evaluation of model performance and the training part for estimating an output nonlinearity. For the LN model, we obtained the spatiotemporal stimulus filter (decomposed into a spatial and a temporal filter as explained above under *Receptive field characterization*) from the spatiotemporal white-noise experiments, but estimated the nonlinearity from the natural-movie data. To do so, we projected the movie frames onto the upsampled spatial filter (to single-pixel resolution) and then convolved the result with the temporal filter. The output nonlinearity was obtained as a histogram (40 bins containing the same number of data points across the range of filtered movie-stimulus signals), containing the average filtered signal and the average corresponding spike count. To apply the nonlinearity to the test data, we used linear interpolation of histogram values. We estimated model performance using the coefficient of determination to obtain the fraction of explained variance (R^2^).

We calculated movie response correlations as the Pearson correlation coefficient between all cell pairs of the same type (as well as across specific pairs of types), using the trial-averaged firing rates of the test stimulus. We performed the same analyses for model predictions and for calculating correlations inherent to the test stimulus, where we calculated pairwise correlations of the light intensity of 5000 randomly selected pixels. To generate correlation-distance curves (**Fig. 1F**), we sorted pairs by ascending distance, and averaged pair correlations over groups of 20-60 pairs (depending on cell type, using fewer pairs per bin when the number of available cells was small).

### Analysis of spatial contrast

To investigate the effects of spatial contrast on response correlations, we split the test (repeated) part of the natural movie into distinct fixations by detecting saccadic transitions. To do so, we first marked each time point when a new image was presented as a transition. Within each image presentation, we calculated the distance between consecutive positions to estimate the instantaneous eye velocity and used MATLAB’s “findpeaks” function to obtain high-velocity transitions. We constrained peak finding for the marmoset (and mouse) to a minimum peak time interval of 47 (and 53) ms and a minimum amplitude of 10 (and 300) deg/s. This process yielded 80 fixations for the marmoset and 68 fixations for the mouse movie.

For each movie frame and each ganglion cell, spatial contrast was calculated as described previously (Karamanlis and Gollisch, 2021) using the standard deviation of pixels inside the cell’s receptive field, weighted by the receptive field weight. For each fixation, we assigned to each cell the median spatial contrast of all frames during the fixation period. We also assigned a linear activation per fixation, estimated by convolving movie frames with the spatial filter obtained from white noise and taking the median over all fixation frames.

To reduce effects of the light level on the analysis of spatial contrast, we aimed at separating the fixations into high-spatial-contrast and low-spatial-contrast groups while balancing the linear activation between the groups. For a pair of cells, we therefore sorted all fixations by the average linear activation across both cells and paired neighboring fixations in this sorted list. This led to 40 pairs (34 for the mouse), and for each pair, we assigned the fixation with the higher spatial contrast to the high-spatial-contrast group and the other fixation to the low-spatial-contrast group. To expand the pairwise correlation (*r_pair_*) into high- and low-spatial-contrast parts, we split the numerator of the Pearson correlation coefficient so that *r_pair_* = *r_high_* + *r_low_*, with

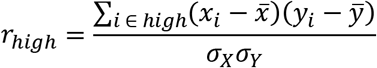

with *x* and *y* corresponding to the responses of the two cells from which the pair was comprised, *i* indexing the frames of the natural movie, and the sum here running over the frames from high-spatial-contrast fixations. Mean 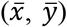 and standard deviation (*σ_X_*, *σ_Y_*) values correspond to the length of the entire movie.

### Extraction of direction-selective (DS) ganglion cells

To identify DS ganglion cells in the mouse retina, we used drifting sinusoidal gratings of 100% contrast, 240 mm spatial period, and a temporal frequency of 0.6 Hz (Sabbah et al., 2017), and analyzed responses as previously described (Karamanlis and Gollisch, 2021). Cells with a mean firing rate of at least 1 Hz and a direction-selectivity-index (DSI) of at least 0.2 (significant at 1% level) were considered putative DS cells. The DSI was defined as the magnitude of the normalized complex sum 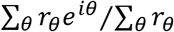. The preferred direction was obtained as the argument of the same sum. The statistical significance of the DSI was determined through a Monte Carlo permutation approach (Karamanlis and Gollisch, 2021; Liu et al., 2017).

To separate ON from ON-OFF DS cells, we used a moving-bar stimulus. The bars (width: 300 μm, length: 1005 μm) had 100% contrast and were moved parallel to the bar orientation in eight different directions with a speed of 1125 μm/s. We extracted a response profile to all bars through singular value decomposition, as previously described (Baden et al., 2016), and calculated an ON-OFF index, to determine whether cells responded only to the bar onset (ON), or to both onset and offset (ON-OFF). Cells with an ON-OFF index (computed as the difference of onset and offset responses, divided by their sum) above 0.4 were assigned as ON DS cells and were grouped into three clusters based on their preferred directions.

### Flashed gratings

Depending on the experiment, we generated 1200 to 2400 different sinusoidal gratings with 25 or 30 different spatial frequencies (*f*), with half-periods between 15 and 1200 μm, approximately logarithmically spaced. For each grating, we generated 12 or 10 equally spaced orientations (*θ*) and 4 or 8 equally spaced spatial phases (*φ_o_*). For a given grating, the contrast value for each pixel with (*x, y*) coordinates were generated based on the following equation:

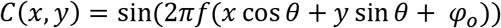

Gratings were presented as 200-ms flashes on the retina, separated by a 600-ms or 800-ms gray screen in between. The order of presentation was pseudorandom. We collected spike-count responses to the flashes by counting spikes during stimulus presentation for the marmoset, or 20 ms after stimulus onset up to 20 ms after stimulus offset for the mouse. We used tuning surfaces to summarize responses (**Fig. S5**), which we generated by averaging responses over trials and spatial phases for each frequency-orientation pair. In the mouse recordings, where we typically collected four to five trials per grating, we calculated symmetrized R^2^ values for the spike counts, and we only used cells with an R^2^ of at least 0.2 for further analyses. In marmoset recordings, we typically collected one to two trials per grating, and we thus used no exclusion criterion.

### Difference-of-Gaussians model

We analytically estimated the activation of a difference-of-Gaussians (DoG) receptive field, by considering the activations of both center and surround elliptical Gaussians to the grating, based on previous calculations (Soodak, 1986). The DoG receptive field was defined with the following parameters: standard deviations *σ_x_* and *σ_y_* at x- and y-axes, the orientation of the x-principal axis *θ_DoG_*, the scaling for the subunit surround *k_s_*, and a factor determining the relative strength of the surround *w_s_*. Concretely, the response of a DoG receptive field (*r_DoG_*) centered at (*x_0_*, *y_0_*), to a parametric sinusoidal grating (*f*, *θ*, *φ_0_*) is

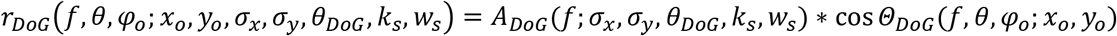

with the amplitude *A_DoG_* given by

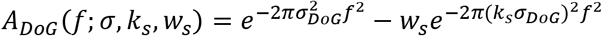

with

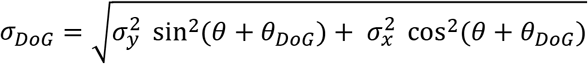

The receptive field phase *Θ_DoG_* is given by

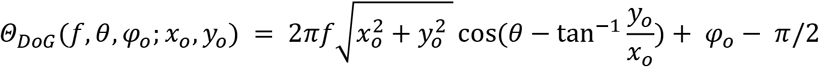

The full DoG model combined the DoG receptive field activation with an output nonlinearity to obtain an LN model of spiking responses. Model response (*R*) was given by

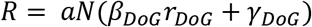

where *N*(*x*) = (1 + *e*^−*x*^)^−1^ is a logistic function and our choice of output nonlinearity, *β_DoG_* and *γ*_DoG_ are parameters determining the steepness and threshold of the output nonlinearity, and *a* is a response scaling factor.

All model parameters (*x_0_, y_0_, σ_x_, σ_y_, θ_DoG_, k_s_, w_s_, β_DoG_, γ_DoG_, a*) were optimized simultaneously by minimizing the negative Poisson log-likelihood, using constrained gradient descent in MATLAB with the following constraints: *σ_x_*, *σ_y_* > 7.5 μm, − *π*/4 < *θ_DoG_* < *π*/4, 1 < *k_s_* < 6, *a* >0. Each trial was used independently for fitting.

### Subunit grid model

We fit all subunit grid models with 1200 potential subunit locations, placed in a hexagonal grid around a given RF center location. The center was taken as the fitted center of the DoG model. The subunits were spaced 16 μm apart. Each subunit had a circular DoG profile, with a standard deviation of *σ* (center Gaussian) and centered at (*x_os_*, *y_os_*), and its activation in response to a grating was given by

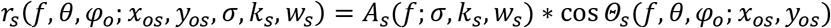

where both amplitude and phase are given by the DoG receptive field formulas with *σ_x_* = *σ_y_* = *σ*.

The full response model was

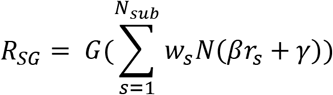

where *N*(*x*) = (1 + *e*^−x^)^−1^ is a logistic function, *β* and *γ* are parameters determining the steepness and threshold of the subunit nonlinearity, *w_s_* are non-negative subunit weights and *G* is a Naka-Rushton output nonlinearity *G*(*x*) = *a x^n^*/(*x^n^* + *k^n^*) + *b*, with non-negative parameters ***θ_out_*** = (*a*, *b*, *n*, *k*).

### Fitting and model selection

We optimized subunit grid models using the stochastic optimization method ADAM (Kingma and Ba, 2014), with the following parameters: batch size =64, β_1_ = 0.9, β_2_ = 0.999, ε = 10^-6^. For the learning rate (η) we used a schedule with a Gaussian profile of μ = Nepochs/2 and σ = Nepochs /5: this led to a learning rate that was low in the beginning of the training, peaked midway, and was lowered again towards the end. Peak learning rate was set to η = 0.005.

To enforce parameter constraints, we used projected gradient descent. The cost function we minimized was

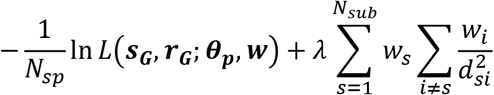

with *N_sp_* being the total number of spikes, *L* the Poisson likelihood, ***s_G_*** the vector of all grating parameters used, ***r_G_*** the corresponding spike-count response vector, ***θ_p_*** = (*σ*, *k_s_*, *w_s_*, *β*, *γ*, ***θ_out_***) all the shared model parameters, and ***w*** = (*w*_1_,…,*w_N__sub_*) the vector containing all subunit weights. *λ* controls the regularization strength, which depends on the pairwise subunit distances *d_si_*. This novel density-based regularizer allowed the coverage of the receptive field with a flexible number of subunits (**Fig. S5**).

After the end of the optimization, we pruned subunit weights with small contributions or weights that ended up outside the receptive field. To do so, we first set to zero every weight smaller than 5% of the maximum subunit weight. We then fitted a two-dimensional Gaussian to an estimate of the receptive field, obtained by summing subunit receptive fields weighted by the subunit weights. The weight corresponding to any subunit center lying more than 2.5 sigma outside that Gaussian was set to zero. To ensure proper scaling of the output nonlinearity after weight pruning, we refitted the output nonlinearity parameters along with a global scaling factor for the weights.

We typically fitted six models per cell with different regularization strengths ranging from 10^-6^ to 5·10^-4^. To select for the appropriate amount of regularization, we only accepted models that yielded at least three subunits and had a low receptive field coverage (less than 3; see below). Amongst the remaining models, we selected the one that minimized the Bayesian Information Criterion (BIC), which we defined for the subunit grid model as

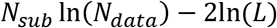

where *N_sub_* is the number of non-zero subunits, *N_data_* is the number of grating-response pairs used to fit the model, and *L* the likelihood of the fitted model. The selected model balanced good prediction performance and realistic receptive field substructure.

### Parameter characterization of the subunit grid model

Subunit coverage summarized how densely subunits covered a cell’s receptive field. A coverage value was calculated if there were at least 3 subunits with non-zero weights in the model. It was calculated as the ratio A/B, where A was the subunit diameter (4σ of the center Gaussian), and B was the average subunit distance. For a particular cell, the average subunit distance was calculated as the average over all nearest-neighbor distances, weighted by each pair’s average subunit weight.

To plot and characterize subunit nonlinearities, we first added an offset so that at an input of zero they show zero output. We then scaled them so that the maximum value over the input range [-1,1] was unity. Following offsetting and scaling, we calculated nonlinearity asymmetries to quantify the response linearity of subunits:

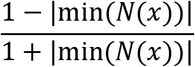

where *x* ∈ [-1,1] and *N*(*x*) is the nonlinearity defined over the same range.

### Natural images and response predictions

We flashes a series of 220 (or 120) natural images to the retina, as described previously (Karamanlis and Gollisch, 2021). We used images from the Van Hateren database, which were cropped to their central 512×512 pixel square, and flashed over the multielectrode array at single-pixel resolution. All images were multiplicatively scaled to have the same mean intensity as the background. Interspersed with the natural images, we also presented artificial images. The images were generated as black-and-white random patterns at a single-pixel level, and then blurred with Gaussians of eight different spatial scales (Schwartz et al., 2012). The images were flashed for 200 ms, interleaved with either 600 or 800 ms of gray background. Images were flashed in a randomized order, and we typically collected eight trials per image. Average spike counts were calculated as in the case of flashed gratings and only cells with symmetrized R^2^ of at least 0.2 were used for further analyses.

To calculate response predictions for models built with white noise, we used the output of spatial filters that were convolved with the natural images. The filters were upsampled to match the resolution of the presented images and normalized with the sum of their absolute values. For models obtained from responses to flashed gratings, DoG receptive fields were instantiated at a single pixel resolution, and the natural images were then projected onto the DoG receptive fields. For the subunit model, the subunit filter outputs were passed through the fitted subunit nonlinearity, and then summed with applying the subunit weights. The performance for each model was calculated as the Spearman’s rank correlation between the model output (without an explicit output nonlinearity) and cell responses to the natural images.

### Flickering gratings and spatiotemporal DoG LN models

We generated 3000 (or 4800) different gratings with 25 (or 30) different spatial frequencies, between 7.5 and 1200 μm half-periods, approximately logarithmically spaced. For each grating, we generated 20 orientations and 6 (or 8) spatial phases. The gratings were presented in a pseudorandom sequence, updated at an 85 (or 75) Hz refresh rate. Every 6120 (or 3600) frames, we interleaved a unique sequence of 1530 (or 1200) frames that was repeated throughout the recording to evaluate response quality.

We fitted a spatio-temporal DoG LN model to the grating responses. The temporal filters spanned a duration of 500 ms and were modeled as a linear combination of ten basis functions. The response delay was accounted for with two square basis functions for each of the two frames before a spike. The remaining eight were chosen from a raised cosine basis (Latimer et al., 2019), with peaks ranging from 0 to 250 ms before a spike.

Concretely, the spatio-temporal DoG model had the form

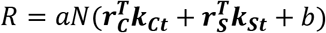

where *N*(*x*) = (1 + *e*^−*x*^)^−1^ is a logistic function, ***k_ct_*** and ***k_st_*** are separate temporal filters for the center and the surround, *b* determines the baseline activation, and *a* is a response scaling factor. The vectors ***r_c_*** and ***r_s_*** contain DoG receptive field activations for 500 ms before a particular frame and were calculated based on the same calculations we used for the flashed gratings. The model was fit with nonlinear constrained optimization, with DoG constraints identical to the case of flashed gratings, and *a* > 0.

### Spatio-temporal subunit grid model

We also fitted a spatio-temporal subunit grid model to the grating responses. Our strategy was very similar to the grating flash case. We fit all subunit grid models with 1200 subunit locations, placed in a hexagonal grid around a given RF center location. The center was taken as the fitted center of the DoG model. The subunits were spaced 16 μm apart. Model response (*R*) was given by

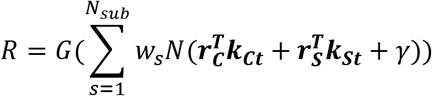

where *w_s_* are non-negative subunit weights, *N*(*x*) is a logistic function, ***k_ct_*** and **k_st_** are separate temporal filters for the center and the surround shared across all subunits, and *γ* determines the nonlinearity threshold. We used a Naka-Rushton output nonlinearity *G*(*x*) = *a x^n^*/(*x^n^* + *k^n^*), with non-negative parameters ***θ_out_*** = (*a, n, k*). The vectors ***r_c_*** (***r_s_***) contain Gaussian center (surround) subunit activations for 500 ms before a particular frame and for each subunit. The parameters required to fit DoG subunits are the standard deviation of the center, the scaling for the subunit surround, and a factor determining the relative strength of the surround.

We used stochastic gradient descent with the ADAM optimizer to fit spatio-temporal models. The parameters where the same as in the flashed-grating models, except for the batch size = 2000 and ηmax = 0.02. As in the case of flashed gratings, we used the same learning schedule for η, and the same regularization to control for subunit density.

### Natural movie predictions of grating-fitted models and maximally-differentiating fixations

To obtain natural movie predictions for models built from flickering gratings, we instantiated either receptive field or subunit center and surround filters to single pixel resolution. Again, we projected movie frames on center and surround filters separately, convolved each result with the corresponding temporal filter, and summed the two outputs for obtaining the final filter output. For subunit grid models, the subunit nonlinearity fitted from the gratings was then applied to the linear subunit outputs, which were then summed with the non-negative pooling weights to obtain the final activation signal. The training part of the movie was used for estimating an output nonlinearity and selecting the regularization strength for the subunit grid model. The output nonlinearity was fitted using maximum likelihood (under Poisson spiking) for both the DoG LN and the subunit grid models and had the same parametric form as the model fit with gratings. Regularization strength was selected based on the log-likelihood of the training set, and eligible models had at least 3 subunits and a receptive field coverage below 3. Model performance was estimated for the test set using the coefficient of determination as a fraction of explained variance (R^2^).

To better differentiate DoG and subunit grid model performance, we selected fixations based on model predictions. For each cell pair, we selected the 20% of the fixations for which the deviation in the prediction of the two models, averaged over the two cells, was largest. For a single cell and a single fixation, the deviations were calculated as the absolute value of model differences normalized by the cell’s overall response range and averaged over all frames of the fixation. Performance of both models (R^2^) was then compared to the frame-by-frame neural response on these fixations and averaged over the two cells. The selection of maximally-differentiating fixations does not favor either model *a priori*, because it is only based on how much model predictions differ and not on their performance in explaining the data.

Similarly to the spatial contrast analysis, we expanded the pairwise correlation (*r_pair_*) into linear and nonlinear contributions, by splitting the numerator of the Pearson correlation coefficient so that *r_pair_* = *r_noniinear_* + *r_linear_*. For a pair of cells, we sorted all fixations (in descending order) by the average deviation of model predictions. We assigned the first half of the fixations to the nonlinear group and the remaining ones to the linear group.

### Contrast-reversing gratings

To compare model fits with classical analyses of spatial integration, we stimulated the retina with full-field square-wave gratings of 100% contrast, as previously described (Karamanlis and Gollisch, 2021). The contrast of the gratings reversed with a temporal frequency of 5 Hz, had 10 spatial frequencies ranging with bar widths ranging from 7.5 to 2000 μm, and, for each spatial frequency, one to eight equidistant spatial phases (with more phases typically assigned to lower spatial frequencies). For all analyses of responses to gratings, we excluded cells with unreliable responses by calculating symmetrized R^2^ values between average response vectors of even and odd trials. We created the response vector of a single trial by concatenating single trial PSTHs from all different spatial frequencies and phases. We only considered cells with R^2^>0.1 for our population analyses.

### Data and code availability statement

Data containing cell responses to natural stimuli and code to fit subunit grid models to grating data will be made available at the time of publication.

## Acknowledgments

We thank Jacob Yates and Jude Mitchell for providing eye movement data from marmosets; Arne Meyer for providing mouse horizontal gaze traces; members of the Gollisch lab for assistance during marmoset data collection; Fred Rieke and Juan Anguera for advice on experiments with the primate retina; Nikoloz Sirmpilatze for comments on the manuscript. This work was funded by the European Research Council under the European Union’s Horizon 2020 research and innovation program (grant agreement number 724822) and by the Deutsche Forschungsgemeinschaft (DFG, German Research Foundation) Project-ID 432680300 (SFB 1456, project B05). D.K. was supported by a Boehringer Ingelheim Fonds fellowship.

## Author contributions

D.K. designed the experiments, collected, and analyzed data with supervision from T.G. M.H.K., H.M.S., S.J.Z., and M.M. assisted with marmoset data collection. S.J.Z. performed spike-triggered NMF analyses. D.K. and T.G. wrote the paper with input from all authors.

---

**Fig. S1.**
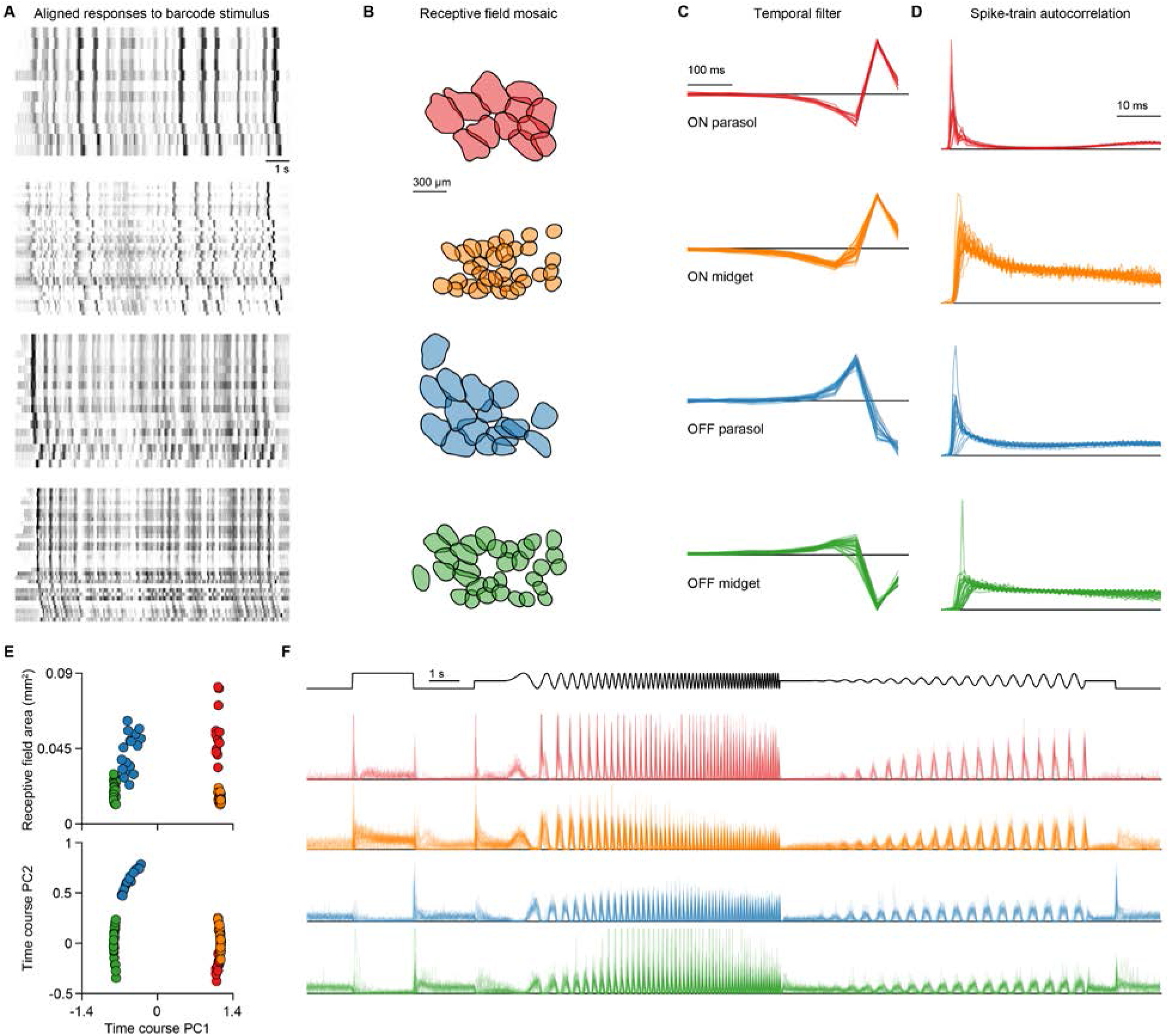
Cell type identification in the marmoset retina. The retina was stimulated with a barcode stimulus. Cell responses were then clustered along with information from white-noise stimulation (receptive field size, temporal filter, autocorrelation). (**A**) Responses to the barcode stimulus of four identified clusters. These responses were aligned to a seed cell (first row) to show the match. (**B**) Receptive-field mosaics of the four identified clusters. (**C**) Temporal filters. (**D**) Spike-train autocorrelations (bin size is 0.5 ms). (**E**) Identified types cluster in simple spaces. Receptive field area was calculated from the estimated contours. (**F**) Responses of the identified cells to a chirp stimulus. PSTHs are calculated with 10-ms bins and normalized to unit sum for plotting.

**Fig. S2.**
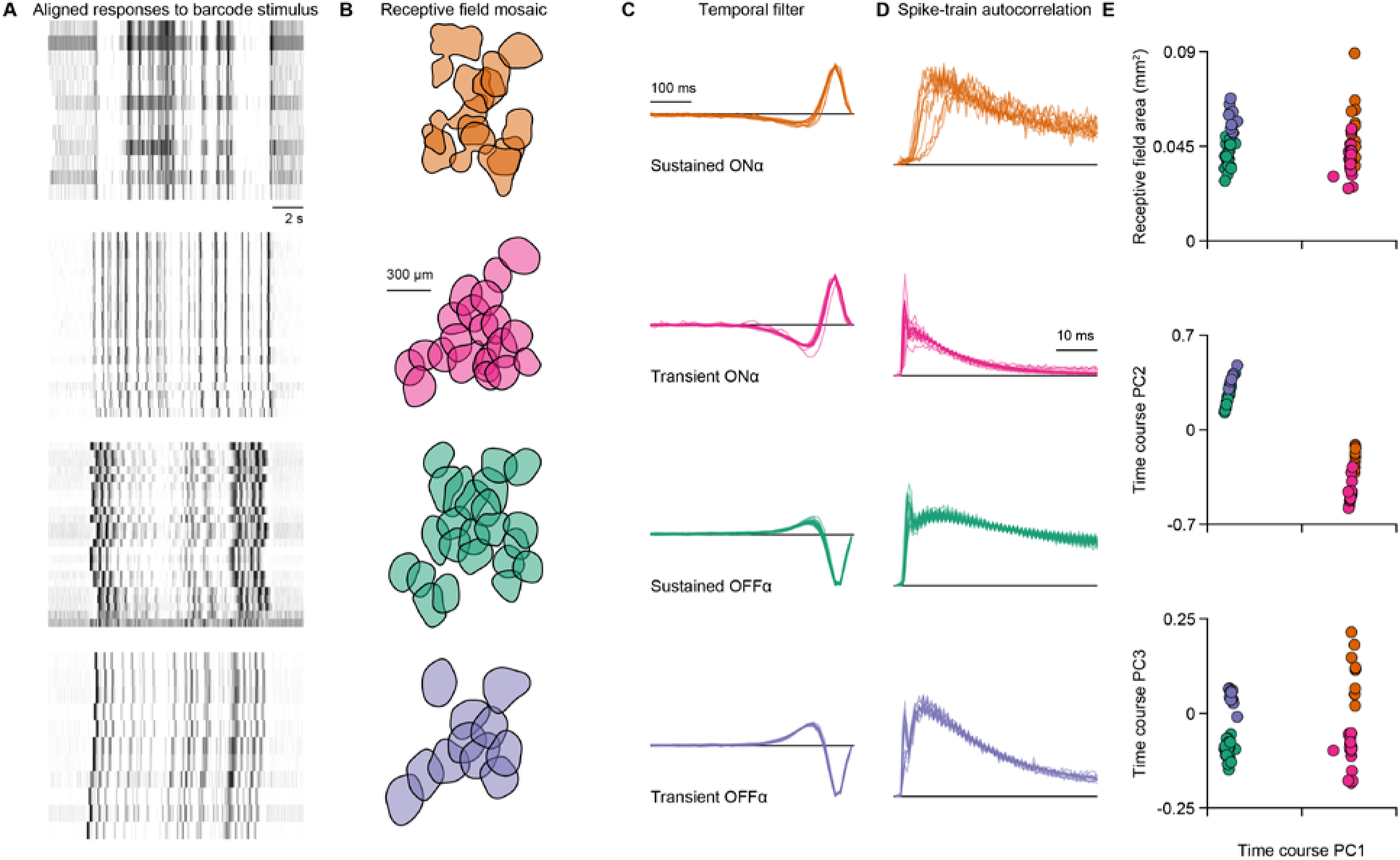
Cell type identification in the mouse retina. The retina was stimulated with a barcode stimulus. Cell responses were then clustered along with information from white-noise stimulation (receptive field size, temporal filter, autocorrelation). (**A**) Responses to the barcode stimulus of four identified clusters. These responses were aligned to a seed cell (first row) to show the match. (**B**) Receptive field mosaics of the four identified clusters. (**C**) Temporal filters. (**D**) Spike-train autocorrelations (bin size is 0.5 ms). (**E**) Identified types cluster in simple spaces.

**Fig. S3.**
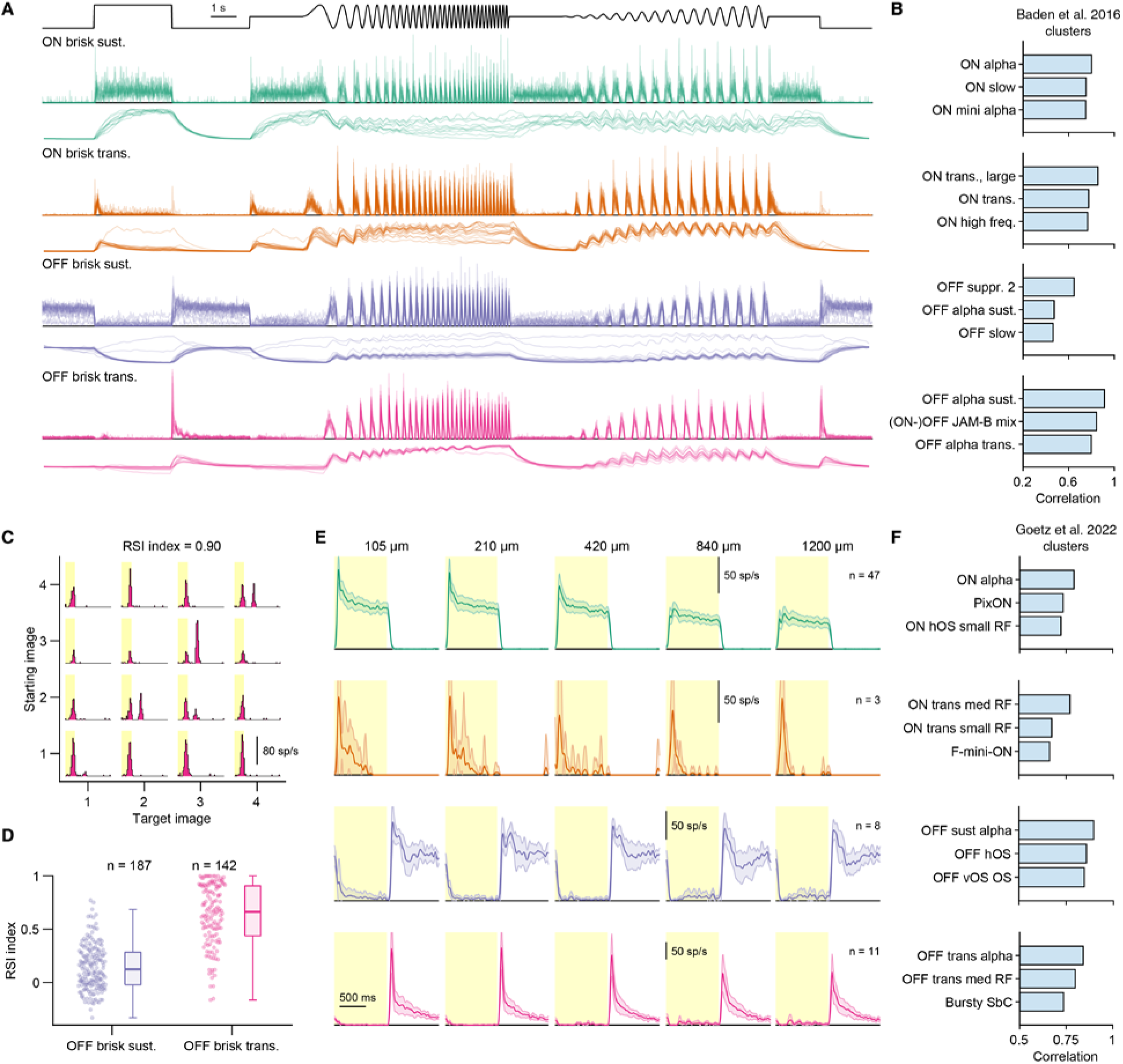
Mapping identified ganglion cell mosaics to alpha types in the mouse retina. (**A**) Responses of four identified ganglion cell types to the chirp stimulus (top), previously used to classify mouse retinal ganglion cells (Baden et al., 2016). The four types are here called ON and OFF brisk sustained and transient cells, according to their response characteristics. The spiking responses were converted to a calcium-equivalent signal (shown below the firing-rate profiles) and (**B**) compared (with Pearson correlation) to the reported templates of different types. We calculated the average correlation of the cells from the four identified types with the reported templates (Baden et al., 2016). Shown are the top three hits. (**C**) An OFF brisk transient cell showing image recurrence sensitivity, measured with saccade gratings (Krishnamoorthy et al., 2017). This sensitivity was quantified with the recurrence sensitivity index (RSI). (**D**) OFF brisk transient cells had significantly higher RSI indices than OFF brisk sustained cells (mean ± SD vs. mean ± SD, mean ± SD, p<10^-50^, Wilcoxon rank-sum test), and the indices were significantly larger than 0.5 (p = 0.038, Wilcoxon sign-rank test), the threshold used for the original characterization. This corroborates that the identified OFF brisk sustained cells correspond to transient-OFFα cells. (**E**) Average responses of the four main types to flashed spots of five different sizes. The spots were flashed either within, or very close to the receptive field centers of the selected cells. Shaded error bars are 95% confidence intervals. (**F**) Spot responses were compared (via correlations) to a functional database (Goetz et al., 2022). Top three hits are shown, generally matching the alpha types. For ON brisk transient cells, the match is ON transient medium RF, hypothesized to match the original description of the transient-ONα cell (Krieger et al., 2017).

**Fig. S4.**
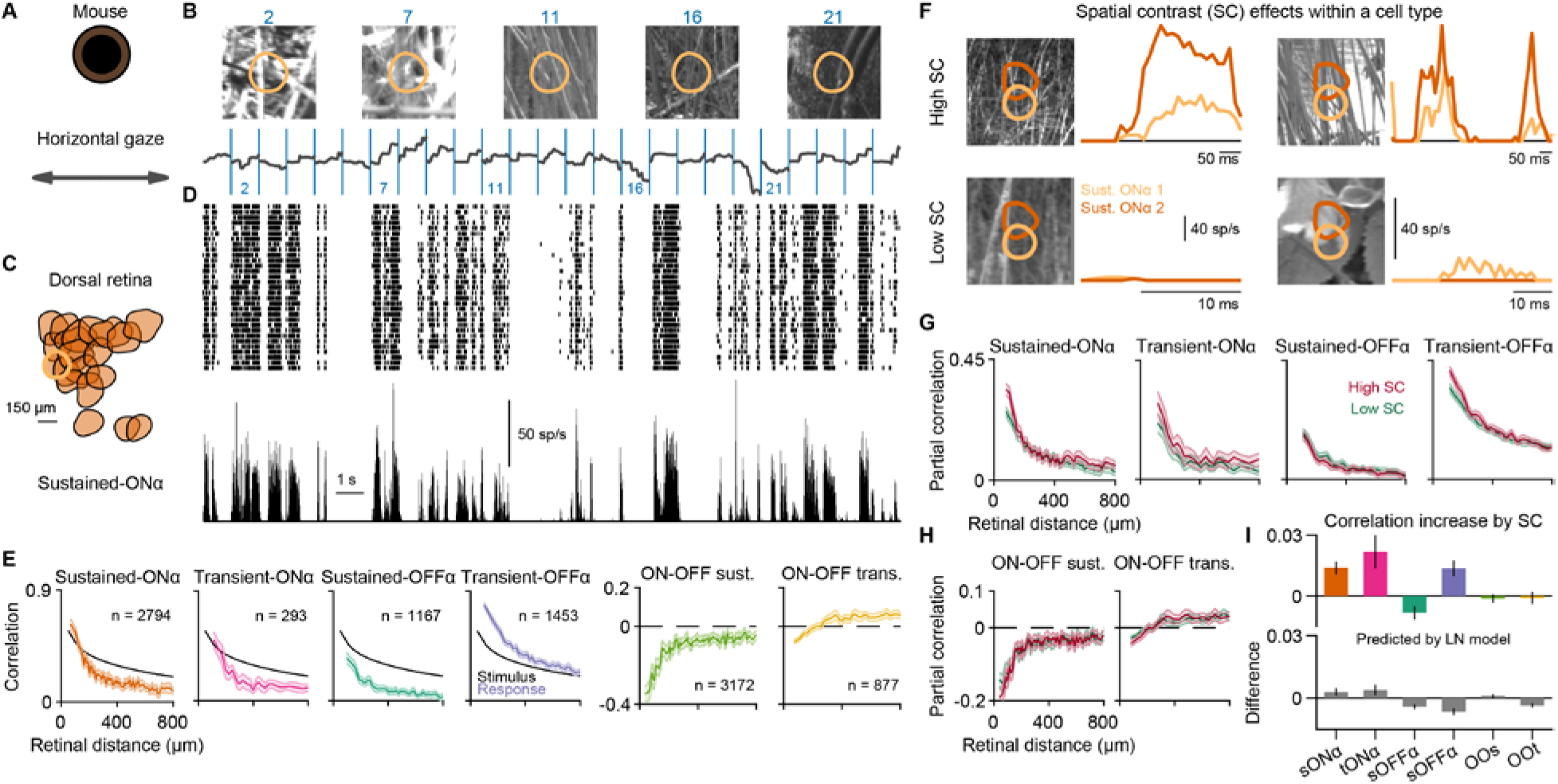
Mouse natural movies, pairwise correlations, and spatial contrast analysis. (**A**) We generated mouse-specific movies by shifting natural images according to horizontal gaze traces recorded from freely-moving mice. (**B**) Each image was presented for 1 s (annotated by the blue lines) and displaced along a cardinal direction that was randomly assigned per image. The receptive field of a sample sustained-ONα cell is overlaid. (**C**) Receptive-field mosaic of simultaneously recorded sustained-ONα cells. (**D**) Spike-raster for 30 trials (top) and the corresponding peri-stimulus-time-histogram (bottom). (**E**) Pearson correlation coefficient between the responses of two ganglion cells as a function of their distance under the natural movie. Colored lines represent average correlation for pairs at similar distance (with 95% confidence interval) within the same ganglion cell type from three retinas. For reference, the correlation between stimulus pixels is shown (black lines). This analysis was also done for pairs of ON and OFF ganglion cells. (**F**) Responses of two neighboring sustained-ONα cells to fixations with similar light intensity, but either high (top), or low spatial contrast. (**G**) For each cell type, pairwise correlations were expanded into a sum of high- and low-spatial-contrast partial correlations. (**H**) Same as (G), but for cell pairs of specific types with different response polarity. (**I**) Median differences between high- and low-spatial-contrast partial correlations varied across types (top), and these differences diverged from differences calculated with classical linear-nonlinear (LN) models fit to cells of the same type (bottom). Error bars are robust confidence interval (95%).

**Fig. S5.**
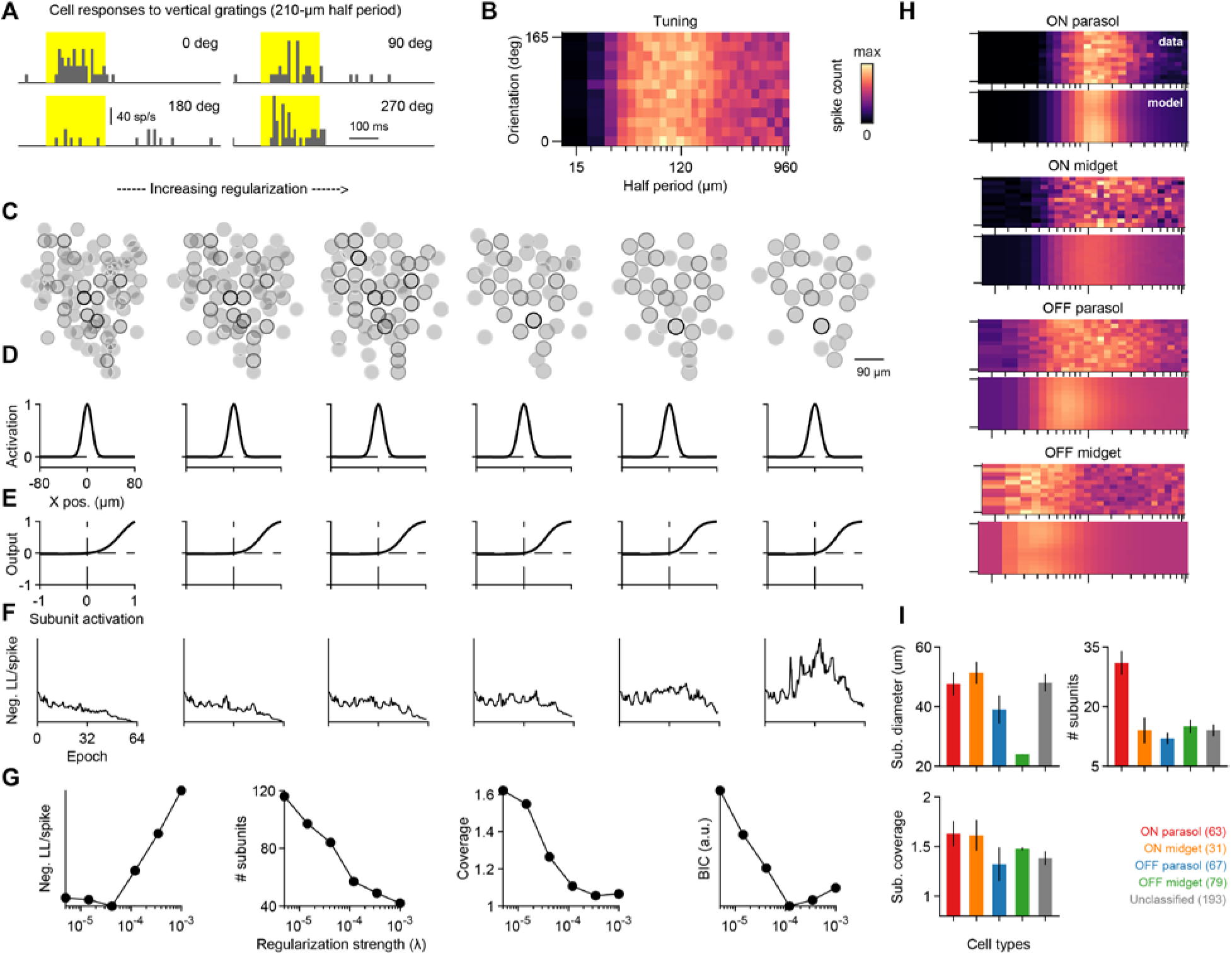
Weight density regularization and ganglion cell responses to flashed gratings. (**A**) Responses of a mouse retinal ganglion cell to flashed gratings of different spatial phases. (**B**) Tuning surface summary of the responses for the same cell, responses for each orientation–spatial period pair were averaged over phases and trials. (**C**) Subunit grids fitted to spiking responses of the sample cell for six different regularization values. Subunit receptive field profiles (**D**) and nonlinearities (**E**) are shown for each fit. (**F**) Parameters were fitted by minimizing the negative Poisson log-likelihood (Neg. LL). Training curves were smoothed with a moving median filter (length of ten points) for plotting. The effects of varying regularization strength (**G**) on the cost function at the end of the optimization, the number of subunits, subunit coverage, and the Bayesian Information Criterion (BIC), which was used to select the best model. We used the number of non-negative subunits as the number of parameters for specifying the BIC. (**H**) All subunit grid models could fit responses to the flashed gratings reasonably well, as demonstrated by the actual tuning surface (top) vs. its prediction from the SG model (bottom), for four sample cells of the marmoset retina (same as **Fig. 2**). (**I**) Subunit diameter, subunit number per cell, and subunit coverage of receptive field for the identified four marmoset ganglion cell types as well as unclassified cells. Error bars are median ± 95% robust confidence interval.

**Fig. S6.**
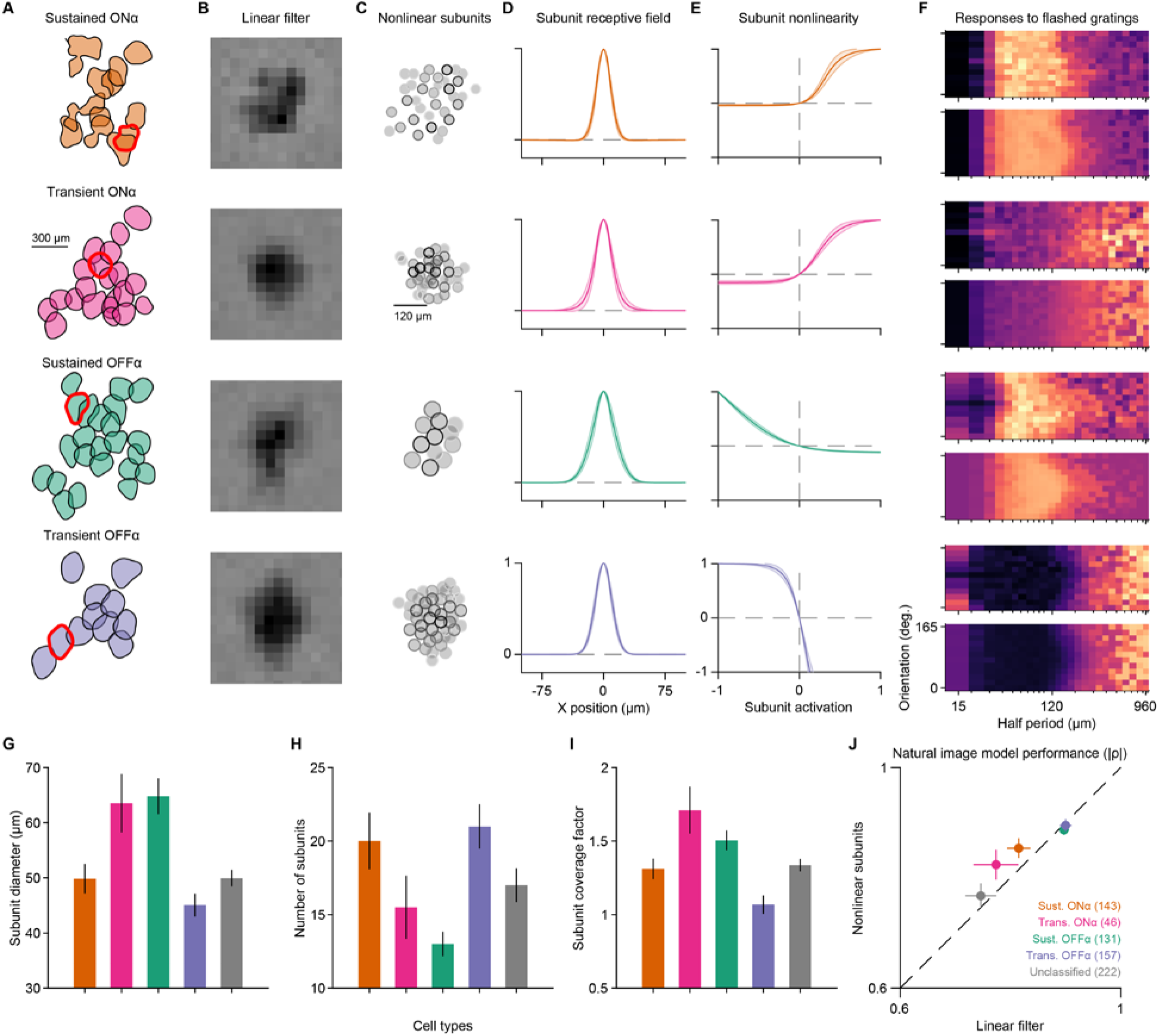
Cell type analysis of model parameters for the mouse retina. We extracted four major cell types from the mouse retina, and the first panels (A-H) depict an example recording with a good representation of all four. (**A**) Receptive-field mosaics for the four types show tiling. Contours were shrunk by 80% for clarity. (**B**) White-noise spatial filter for the highlighted cell in each mosaic. (**C**) The subunit grid map for the highlighted sample cells. (**D**) Subunit profiles for the cells in the mosaic. Shaded areas show 95% robust confidence intervals around the mean. (**E**) Subunit nonlinearities of the fitted models. (**F**) All subunit grid models could fit responses reasonably well, as demonstrated by the actual tuning surface (top) vs. its prediction from the SG model (bottom), shown here for the four sample cells. (**G**) Median subunit diameters for all four types. (**H**) Median subunit numbers for all types. (**I**) Median coverage of the subunit mosaic. (**J**) Performances of the subunit grid model (“nonlinear subunits”) and the LN model (“linear filter”) in predicting responses to natural images, calculated as the absolute value of the Spearman’s ρ between predictions and measured average spike counts. Error bars in (G)-(J) are median ± 95% robust confidence interval.

**Fig. S7.**
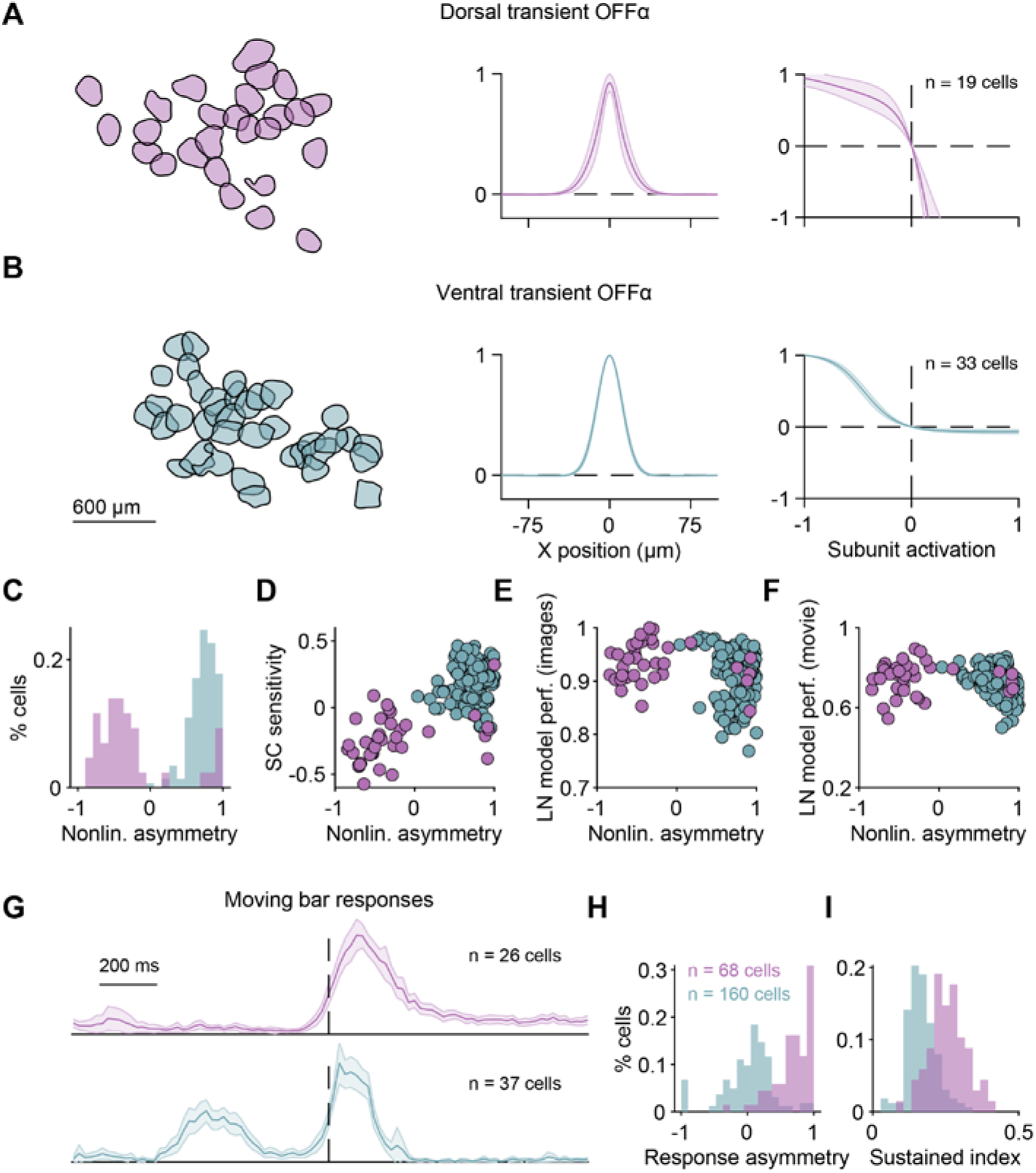
Subunit nonlinearities differ between dorsal and ventral transient-OFFα cells. (**A**) Subunit model parameters for a transient-OFFα cell mosaic in the dorsal retina. (**B**) Same as (A), but for a recording from the same eye coming from the ventral retina. (**C**) Asymmetry in the nonlinearities is evident for all recorded transient-OFFα cells. (**D**) The asymmetry affects sensitivity to spatial contrast (SC) in natural scenes. Spatial-contrast sensitivity was measured as described previously (Karamanlis and Gollisch, 2021). Nonlinearity asymmetry was related to both model performance calculated for natural images (**E**) but also natural movies (**F**). Both dorsal and ventral cells were better predicted with LN models if their nonlinearity asymmetries were close to zero. (**G**) Firing rate responses (normalized) of dorsal (top) and ventral (bottom) transient-OFFα cells to a moving bar stimulus. The responses correspond to the average bar response over eight different directions. The bars had an ON contrast and approximately entered the receptive field of the cells at the start of the displayed traces and left the receptive field approximately at the timepoint marked by the dashed lines. The responses in the ventral retina showed a peak following the onset of the bar, which we quantified with a response asymmetry index (**H**). The index was defined as (Roff - Ron)/(Roff + Ron), where Ron and Roff are the average responses before and after the bar leaves the receptive field center. This index was significantly larger for the dorsal retina (0.67 ± 0.31 vs 0.00 ± 0.38, mean ± SD, p < 10^-23^, Wilcoxon rank-sum test). (**I**) Moving bar offset responses in the dorsal retina were more sustained compared to the ventral retina (0.26 ± 0.06 vs 0.23 ± 0.22, mean ± SD, p < 10^-12^, Wilcoxon rank-sum test). The sustained index was defined as the ratio of the average response over the maximum response in the time window following the bar leaving the receptive field center.

**Fig. S8.**
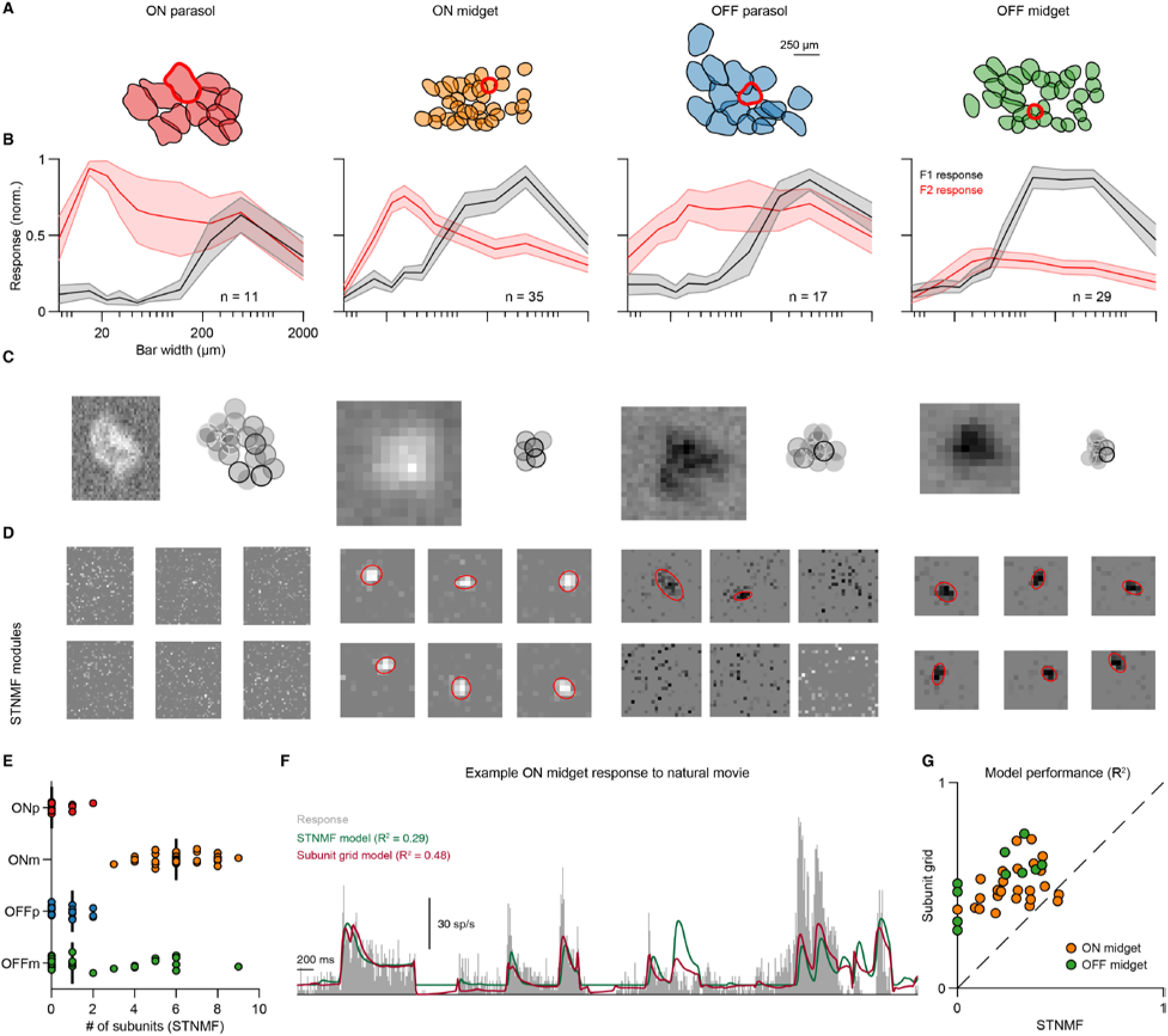
Method comparison with spike-triggered non-negative matrix factorization (STNMF). (**A**) Receptive-field mosaics from a peripheral marmoset retina recording. Sample cells are marked with red outlines. (**B**) Summary of responses to contrast-reversing gratings of different spatial frequencies. Spatial frequency tuning curves for the first Fourier harmonic (F1; black) and the second Fourier harmonic (F2; red). F1 is calculated as the maximum and F2 as the mean harmonic of the responses of the cells over all spatial phases. The error bars represent the SEM. For all four types, the effect of a suppressive surround is clear as both F1 and F2 components decay with increasing bar width. Except for OFF midget cells, the spatial nonlinearity is evident in the strong F2 component for small stimulus scales. (**C**) Spatial filter of a sample cell and the corresponding subunit grid fitted with flickering gratings. (**D**) STNMF applied to a one-hour-long recording of spatiotemporal white-noise stimulation with high spatial resolution. STNMF recovers subunits for midget cells, but here fails for parasol cells, likely owing to the large number of pixels that need to be included for these cells. (**E**) Number of subunits recovered by STNMF for cells of all four types. Vertical black lines mark the medians. (**F**) Using responses to the natural movie, we compared the prediction performance of the subunit grid model and a subunit model derived from STNMF. For the latter model, STNMF subunit outputs were rectified and summed to obtain generator signals. Summation weights were determined by fitting a linear combination of subunit filters to obtain the overall spatial filter, using non-negative least squares. Generator signals were then related to spiking responses by fitting a logistic output nonlinearity using the non-repeated part of the natural movie (as for the subunit grid model). (**G**) Model performance comparison for midget cells between the two nonlinear subunit methods. Parasol cells were omitted because STNMF failed to recover meaningful decompositions.

**Fig. S9.**
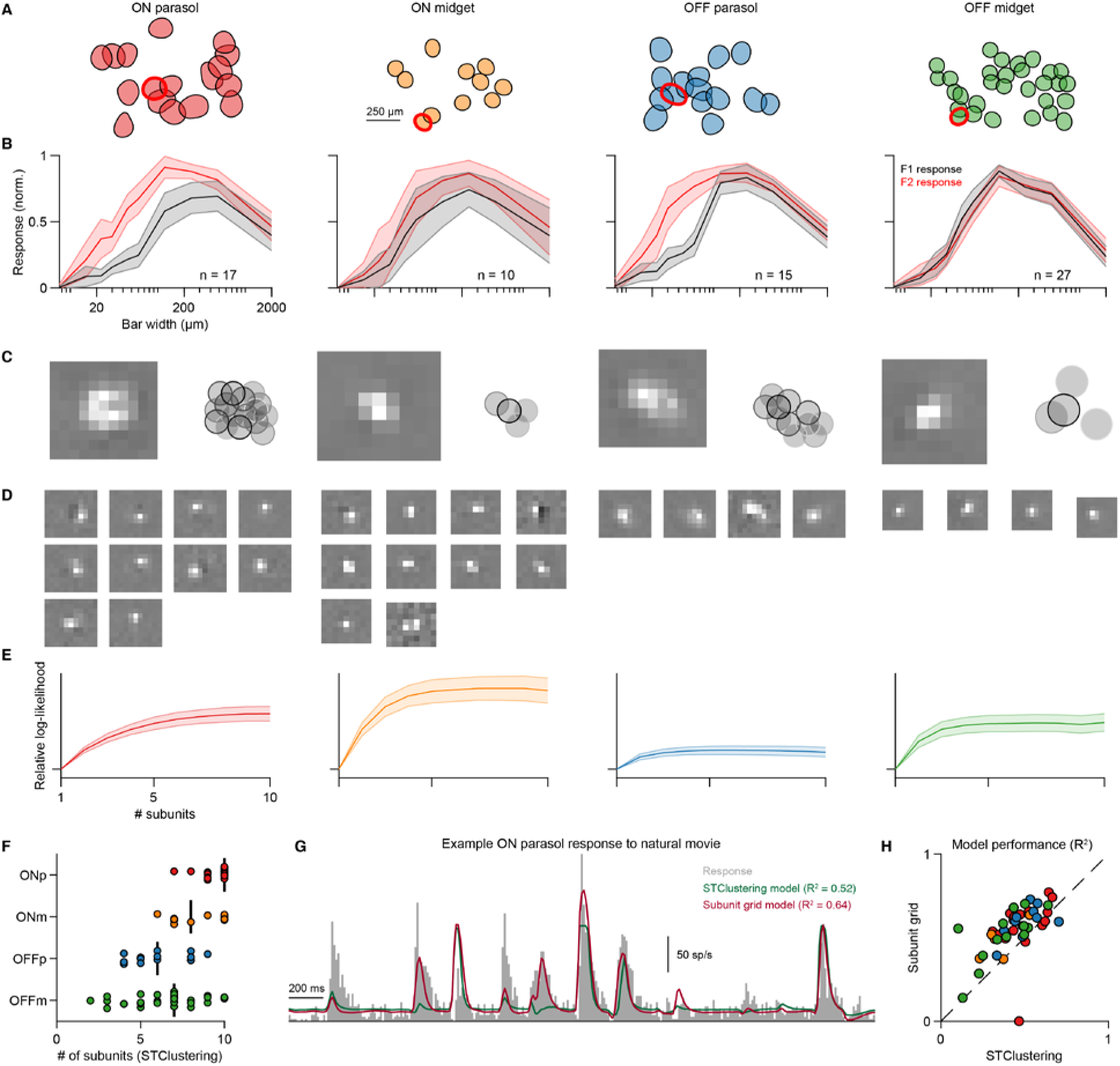
Method comparison with spike-triggered clustering (STClus). (**A**) Receptive-field mosaics from a peripheral marmoset retina recording. Sample cells are marked with red outlines. (**B**) Summary of cell type responses to contrast-reversing gratings, shown as in **Fig. S8B**. (**C**) The receptive field of a sample cell (left) and the corresponding nonlinear subunit decomposition obtained by the subunit grid method. (**D**) Nonlinear subunit decomposition obtained by STClus (Shah et al., 2020) applied to white-noise responses. The number of subunits selected maximized the likelihood of a validation set. (**E**) The log-likelihood for different numbers of subunits for all cells of the same type. Error bars are 95% confidence intervals. (**F**) Number of subunits that maximized the validation likelihood for each cell. Black bars are medians over cells belonging to the same type. (**G**) We compared the prediction performance of the two models using the natural movie. STClus subunit outputs were exponentiated and summed to obtain generator signals. Summation weights were determined by the STClus fitting procedure using the white-noise data. Generator signals were then related to spiking responses by fitting a model-specific output nonlinearity (Shah et al., 2020), using the non-repeated part of the natural movie (as for the subunit grid model). (**H**) Model performance comparison between the two nonlinear subunit methods.

**Fig. S10.**
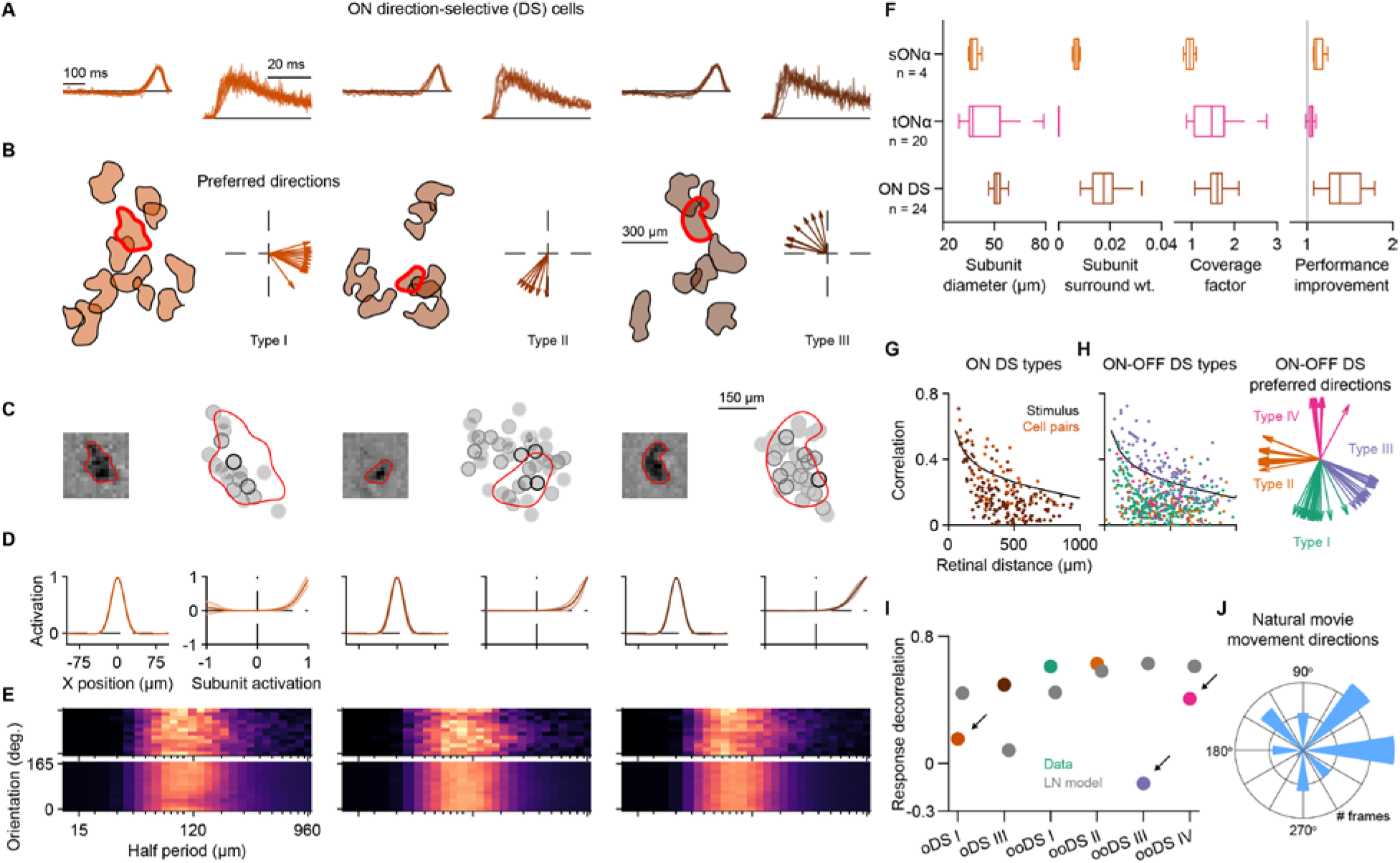
Direction-selective cells show correlated responses when driven by natural movies. (**A**) ON direction-selective (DS) cells have monophasic filters, with autocorrelograms suggesting sustained spiking responses. (**B**) ON DS cells were split into three subtypes, based on their preferred directions under a drifting grating stimulus. (**C**) Receptive fields and subunit layouts obtained from the subunit grid model for three sample ON DS cells. Note that the subunit map only roughly matches the receptive-field contour. (**D**) ON DS cells have small subunits with strong rectification. Shaded error bars depict 95% confidence intervals. (**E**) The tuning surfaces of ON DS cells revealed strong suppression for higher spatial scales. (**F**) We compared ON DS model fits with sustained-(sONα) and transient-ONα cells (tONα) from the same recording. Compared to the other two ON types, ON DS cells had larger subunit diameters, stronger subunit surround, comparable coverage factors, and larger performance improvement for natural images. Numbers denote the number of cells in each group. (**G**) Pairwise correlations for ON DS cells from two subtypes (I and III) under natural movie stimulation. Cells from subtype II were excluded because they had unreliable responses to the movie. Each spot corresponds to a pair. (**H**) Same as (G) but for ON-OFF DS cell pairs (left). ON-OFF DS cells were clustered based on their preferred directions (right). (**I**) A subtype from ON DS cells (I) and two from ON-OFF DS (III and IV) showed low stimulus decorrelation (colored circles, arrows), lower than what would be predicted by an LN model (gray circles). (**J**) Large gaze shifts (>75μm/frame) in our constructed natural movie typically caused global movement with a strong rightward component, approximately matching the preferred directions of cells with relatively low decorrelation values. We hypothesize that this prevalence of motion in the preferred direction led to the increased concerted activity of these DS cells.

